# Exosome-mediated MIR211 modulates tumor microenvironment via the DUSP6-ERK5 axis and contributes to BRAFV600E inhibitor resistance in melanoma

**DOI:** 10.1101/548818

**Authors:** Bongyong Lee, Anupama Sahoo, Junko Sawada, John Marchica, Sanjay Sahoo, Fabiana I. A. L. Layng, Darren Finlay, Joseph Mazar, Piyush Joshi, Masanobu Komatsu, Kristiina Vuori, Garth Powis, Petrus R. de Jong, Animesh Ray, Ranjan J. Perera

## Abstract

The microRNA MIR211 is an important regulator of melanoma tumor cell behavior. Previous studies suggested that in certain tumors, MIR211 acted as a tumor suppressor while in others it behaved as an oncogenic regulator. When MIR211 is expressed in BRAFV600E-mutant A375 melanoma cells in mouse xenografts, it promotes aggressive tumor growth accompanied by increased cellular proliferation and angiogenesis. We demonstrate that MIR211 is transferred to adjacent cells in the tumor micro-environment via exosomes. Cross-species genome-wide transcriptomic analysis showed that human tumor-derived MIR211 interacts with the mouse transcriptome in the tumor microenvironment, and activates ERK5 signaling in human tumor cells via the modulation of a feedback loop. Human miR211 directly inhibits human DUSP6 protein phosphatase at the post-transcriptional level. We provide support for the hypothesis that DUSP6 inhibition conferred resistance of the human tumor cells to the BRAF inhibitor vemurafenib and to the MEK inhibitor cobimetinib, with associated increases in ERK5 phosphorylation. These findings are consistent with a model in which MIR211 regulates melanoma tumor proliferation and BRAF inhibitor resistance by inducing ERK5 signaling within the complex tumor microenvironment. We propose that the MIR211-ERK5 axis represents an important and sensitive regulatory arm in melanoma with potential theranostic applications.

## INTRODUCTION

Malignant melanoma has a high mortality burden if not detected early(1,2). The histopathological differentiation of benign pigmented lesions (melanocytic and Spitz nevi) and malignant melanoma often presents a diagnostic challenge, driving recent efforts to develop novel biomarkers(3,4). Moreover, there continues to be a need to identify molecular master regulators and checkpoints to target with novel drugs. Over the past decade, micro-(mi)RNAs have emerged as promising diagnostic, therapeutic, and theranostic candidates due to their emerging roles in melanoma hallmarks including proliferation, migration, apoptosis, immune responses, and in shaping the tumor niche(4,5).

miRNAs are noncoding 21-23 nucleotide molecules that mostly bind to mRNA 3’UTRs to degrade their target and inhibit protein translation(6). miRNAs are critical regulators of tumor growth, invasion, angiogenesis, and immune evasion through their control of target gene expression(7,8), not least in melanoma(9–14). We discovered a significant role for MIR211 in melanocyte and melanoma pathobiology(15,16), with subsequent studies confirming MIR211 as a critical regulatory molecule in melanoma(17–22).

We previously showed that MIR211 is highly expressed in primary melanocytes; expression is low or absent in 70% of melanomas, and that expression is reduced in melanotic and amelanotic melanoma cell lines(15). Others have reported decreased MIR211 expression in clinical melanoma samples (23,24), driving efforts to use MIR211 as a clinical diagnostic test to discriminate melanomas from benign nevi(25). We also showed that MIR211 overexpression increases mitochondrial respiration in amelanotic melanoma cells by inhibiting pyruvate dehydrogenase kinase 4 (PDK4), thereby inhibiting melanoma cell survival and invasion(16).

Despite a body of data suggesting a tumor suppressor role for MIR211, *Guar et al*.(26) reported that MIR211 is overexpressed in a majority (6/8) of melanoma lines in the NCI-60 cancer cell line panel. MIR211 was also overexpressed in 9/29 melanomas previously analyzed by us(15). Recent studies suggest that: (i) MIR211 is transported via melanosomes to the tumor microenvironment to activate MAPK signaling and thus to promote melanoma growth(11); (ii) MIR211 expression and the melanin biosynthetic pathways are induced by vemurafenib (a BRAF inhibitor) treatment and to contribute to vemurafenib resistance(27); and (iii) MIR211 contributes to BRAF inhibitor resistance in melanoma via MAPK signaling(28). Therefore, although MIR211 clearly participates in melanomagenesis, there is considerable complexity (tumor-promoter or suppressor) in its mechanisms of action that requires further clarification.

Here we interrogated the molecular action of MIR211 *in vivo*. MIR211 overexpression in *BRAF*^V600E^-mutant A375 melanoma cells promoted aggressive xenografted tumor growth, accompanied by cell proliferation and angiogenesis, without marked disruption of the glycolysis or tricarboxylic (TCA) pathways, suggesting that MIR211 drives cancer via at least one other pathway independent of PDK4. Interrogation of xenograft RNA-seq data revealed activation of the ERK5 pathway, which is negatively regulated by the putative MIR211 target gene products BIRC2 and DUSP family members. These genes were confirmed as direct MIR211 targets by RNA immunopurification with anti-Argonaute-2 antibodies and RNA-seq (RIP-seq) and target cleavage assays. Finally, MIR211 conferred resistance to the BRAF inhibitor vemurafenib and MEK inhibitor cobimetinib, with corresponding increases in ERK5 phosphorylation. Taken together, these observations are consistent with a model in which MIR211 modulates melanoma tumor proliferation and resistance to drugs that target the BRAF pathway by inducing signal transduction through the ERK5 pathway.

## MATERIAL AND METHODS

### Cell culture

Melanoma cell lines, A375 (RRID:CVCL_0132) and SKMEL28 (RRID:CVCL_0526) were obtained from the ATCC (American Type Culture Collection, Manassas, VA). LOX-IMV1 (RRID:CVCL_1381) was purchased from the National Cancer Institute (NCI). 501 Mel and A375 derived vemurafenib resistant cell lines(27), A375-P1, A375-P2, A375-C2, and A375-C3, were generously provided by Dr. Laura Poliseno at the Institute of Clinical Physiology (Pisa, Italy). All cells were grown in complete Tu medium as previously described(29). For vemurafenib resistant cells, 2 uM of vemurafenib (PLX4032, Selleckchem, Houston, TX) was added. The cells were grown in a humidified incubator at 37°C, 5% CO_2_.

### Immunohistochemistry

Formalin-fixed paraffin embedded tissues were stained with routine H&E or used for immunohistochemistry. The slides were placed on the Leica Bond RX and stained using primary antibody. Heat retrieval was utilized using epitope retrieval solution 1 (Citrate buffer pH6) for 20 minutes. Antibody was incubated at room temperature for 1hr and was visualized by using Leica’s Bond Polymer Refine detection kit (DS9800). Slides were counterstained with hematoxylin from the kit then slides were dehydrated and coverslipped. The following primary antibodies were used: anti-Ki67 (Thermo Fisher Scientific, Waltham, MA; MA5-14520; 1:100), anti-CD31 (Dianova, Barcelona, Spain; Clone SZ31 P/N DIA-310; 1:25), and anti-PDK4 (LSBio, Seattle, WA; LS-B3459; 1:200). For visualizing CD31, Leica’s Bond Polymer Refine Red detection kit (DS9390) and DAPI (Invitrogen, Carlsbad, CA; D1306 diluted at 1:100 for 5 minutes at RT) were used. Slides were coverslipped using Prolong Gold Antifade Mountant (P/N P36930).

For hypoxy region detection, mice were injected intravenously with 60 mg/kg of pimonidazole (Hypoxyprobe-1 Plus kit, HPI Inc., Burlington, MA) before sacrificing mice. Formalin fixed paraffin embedded samples where cut at 5uM and allowed to air dry. The slides were placed on the Leica Bond RX and stained using supplied primary and secondary antibodies.

All slides that were bright field chromogenic IHC were scanned using the Aperio Scanscope XT system (Aperio Technologies, Inc., Vista, CA) at 20x resolution and the fluorescent scans were from the Aperio Scanscope FL (Aperio Technologies) at 20x resolution.

### Luciferase reporter gene assays

The 3’UTR of DUSP3, 4, 6, and 19 and BIRC2 was amplified using specific primers (**Supplementary File 1**) and cloned downstream of luciferase gene into pcDNA6/Luc(30) using an In-Fusion HD cloning kit (Takara Bio USA, Inc., Mountain View, CA). The recombinant pcDNA6/LUC-DUSP (or pcDNA6/LUC-DUSP-211Δ), pcDNA4/MIR211(31), and pRL-CMV (a *Renilla* luciferase plasmid, Promega, Madison, WI) plasmids were transfected into HEK293T cells (Thermo Fisher Scientific) using FuGENE HD (Promega, Madison, WI). For BIRC2 reporter assay, pcDNA6/LUC-BIRC2 or pcDNA6/LUC-BIRC2-211ΔΔ plasmids were used. At 48 h post-transfection, both firefly and renilla luciferase activities were measured by the Dual-Glo luciferase Assay (Promega) using a GloMax 20/20 Luminometer (Promega). Firefly luciferase activities were normalized with renilla luciferase activities.

### siRNA-mediated knockdown

siRNA targeting ERK5 (catalog no. 4427038, ID: s11149, and catalog no. 4427037, ID: s11150), DUSP3 (catalog no. 4427038, ID: s4371) DUSP4 (catalog no. 4427037, ID: s4372), DUSP6 (catalog no. 4427037, ID: s4378), DUSP19 (catalog no. 4427038, ID: s44472) and BIRC2 (catalog no. 4427038, ID: s1450) were purchased from Ambion (Foster City, CA). siRNAs were transfected at 20 nM for 48h using Lipofectamine RNAiMAX (Life Technologies, Carlsbad, CA). The efficiency was determined by qRT-PCR using TaqMan assay.

### 2D and 3D cellular growth assays

For growth inhibition assay by XMD8-92 (Selleckchem, Houston, TX), cells were seeded at a density of 4000 cells per well in a 96-well plate and compounds were added the next day. Cell viability was determined after 6 days. For growth inhibition assays with AX15836 (Tocris, Minneapolis, MN) in 2D culture systems, cells were seeded at a density of 1000 cells/well in 384-well plates (3 biological replicates). Compounds were added subsequent to cell seeding and diluted twofold to create eighteen-point titrations. Control cells were treated with DMSO. Cell viability was assessed after 5 days by XTT Cell Proliferation Assay (Biotium, Fremont, CA) according to the manufacturer’s instructions.

For growth inhibition assays with AX15836 in 3D cell cultures, cells were seeded at a density of 5000 cells/well in 96-well plates (3 biological replicates) in soft agar. Compounds were added 2 days after cell seeding and diluted twofold to create eight-point titrations. Control cells were treated with dimethylsulfoxide (DMSO). Cell viability was assessed after 10 days by Alamar Blue Cell Proliferation Assay (Thermo Fisher Scientific) according to the manufacturer's instructions.

For growth inhibition assays with vemurafenib (PLX4032, Selleckchem, Houston, TX) in 2D cell cultures, cells were seeded at a density of 1000 cells/well in 384-well plates (3 biological replicates). Compound was added subsequent to cell seeding and diluted twofold to create an eighteen-point titration. Control cells were treated with DMSO. Cell viability was assessed after 5 days by XTT Cell Proliferation Assay according to the manufacturer’s instructions.

### Western blotting

Whole cell extracts were fractionated by SDS-PAGE and transferred to a polyvinylidene difluoride membrane. After incubation with 5% bovine serum albumin (Catalog no. BP1600-100, Fisher Scientific, Hampton, NH) in TBST (10 mM Tris, pH 8.0, 150 mM NaCl, 0.5% Tween 20) for 60 min, the membrane was washed with TBST and incubated with antibodies against phospho-ERK5 (EMD Millipore, catalog no. 07-507), ERK5 (Cell Signaling, catalog no. 3552), phospho-ERK1/2 (Cell Signaling, catalog no. 4370), ERK1/2 (Cell Signaling, catalog no. 4695), and GAPDH (Santa Cruz Biotechnology) at 4°C for overnight. Membranes were washed and incubated with a 1:5000 dilution of horseradish peroxidase-conjugated anti-mouse or anti-rabbit antibodies for 1 h. Blots were washed and developed with the ECL system (Amersham Biosciences, Piscataway, NJ) according to the manufacturer’s protocols.

### RNA immunopurification (RIP) in A375/211 cells

RIP assays in A375/211 cell extracts were performed as described previously (32,33). Briefly, cells were resuspended in lysis buffer (150 mM KCl, 25 mM Tris/Cl pH 7.4, 5 mM EDTA, 0.5% Nonidet P-40, 5 mM DTT, 1 mM PMSF, 1x protease inhibitor cocktail (Pierce, Thermo Fisher Scientific), 100 U/ml SUPERaseIn (Ambion, Austin, TX)). Lysates were spun, and the supernatant was incubated with Dynal My1 streptavidin-coated magnetic beads (Invitrogen, Carlsbad, CA) coupled to biotinylated AGO-specific 4F9 antibody (Santa Cruz, catalog no. sc-53521). The beads were washed twice with ice-cold lysis buffer and RNA was extracted with TRIzol and the miRNeasy RNA extraction kit (Qiagen, Valencia, CA) and stored at −80°C.

### RNA-Sequencing

RIPed RNAs were subjected to double strand cDNA synthesis using an Ovation RNA-Seq System V2 (NuGen Technologies, Inc., San Carlos, CA). High-throughput sequencing libraries were constructed using the TruSeq ChIP Sample preparation kit (Illumina, San Diego, CA) and sequenced on a HiSeq 2500 (Illumina) with 50 bp read lengths. The enriched peaks were defined by MACS (43). For xenografts RNA-seq, total RNA was used for constructing libraries using the Illumina TruSeq Stranded Total RNA Library preparation kit (Illumina Inc). Single-end 50 base-pair sequencing was performed on Illumina HiSeq 2500.

### Mouse xenografts

Severe combined immunodeficiency (SCID) mice (five weeks old) were injected subcutaneously with 5.0 x 10^6^ cells/site (n = 5 per group) into lower left flank. Tumor size was monitored with digital calipers and tumor volume in mm^3^ was calculated by the formula: volume = (width)^2^ x length/2. Mice were euthanized and weights and pictures of excised tumors were taken.

### RNA extraction and real-time quantitative reverse transcription-PCR (qRT-PCR)

Total RNA was purified using the Direct-zol RNA Miniprep kit (Zymo Research, Irvine, CA). For RNAs from xenografts, tumor tissues were pulverized and then used for purification. Quantitative PCR was carried out using TaqMan miRNA assays or SYBR Green mRNA assays as previously described(34). Primer sequences are listed in **Supplementary File 1**.

### Exosome purification and qRT-PCR

Exosomes were purified using ExoCap™ Exosome Isolation and Enrichment Kits (MBL International Corp., Woburn, MA) according to manufacturer’s protocol. Briefly, cells were plated and after 24 h, media were replaced with media containing exosome-depleted fetal bovine serum (FBS; Thermo Fisher Scientific, Waltham, MA) for 48 h. The conditioned media were collected, spun, and filtered to remove cells. The filtered conditioned media were used for exosome purification. Nucleic acids were isolated from captured exosomes in 10 ul volume. One ul out of 10 ul RNA was used for cDNA synthesis in a 15 ul reaction volume using a TaqMan™ MicroRNA Reverse Transcription Kit (Thermo Fisher Scientific). qRT-PCR was performed in a 20 ul reaction volume with 1 ul of cDNA. The cycle threshold (Ct) value was used to compare the levels of miRNAs.

### Targeted metabolomics analysis

Targeted metabolomics of cells and xenografts was performed at the Cancer Metabolism Core at Sanford Burnham Prebys Medical Research Institute (SBPMRI; La Jolla, CA, USA) as previously described(35). Briefly, tumor samples (15-30 mg) were transferred to 2 ml round-bottom tubes (Qiagen, Hilden, Germany) with the addition of a 5 mm stainless steel bead (Qiagen) and ice-cold 50% methanol/20 μM L-norvaline (450 μl). Tubes were shaken at maximum speed (30) for 2 min (Qiagen Tissuelyser), vortexed, and placed on dry ice for 30 min. After thawing on ice, samples were centrifuged (15,000 x g, 10 min, 4°C). The supernatant was transferred to a new tube, vortexed with chloroform (220 μl), and centrifuged (15,000 x g, 15 min, 4°C). The top layer was dried (MiVac, 4 h), derivatized, and analyzed by GC-MS to quantify small polar metabolites as previously described(35).

### Melanoma Cell lines

Drug response sensitivities for vemurafenib in 39 melanoma cell lines have been described previously ^a,b,c^, and the area under the curve (AUC) data were utilized in Supplemental Table 6, to correlate MIR211 expression with vemurafenib responsiveness.

### Prediction of putative MIR211 targets

microRNA.org (http://www.microrna.org) and miRDB (http://www.mirdb.org) were used for target gene prediction as previously described (36,37). Targets overlapping between the two web sites were selected.

### Ingenuity pathway analysis (IPA)

To analyze pathways affected by MIR211, differentially expressed genes between A375 and A375/211 xenografts were compiled and analyzed using Qiagen’s IPA. Analysis was conducted via the IPA web portal (www.ingenuity.com).

## RESULTS AND DISCUSSION

### Ectopic MIR211 expression promotes *in vivo* tumor growth

The amelanotic melanoma cell line A375, which has a constitutively activated MAPK pathway due to *BRAF*^V600E^ mutation and low basal MIR211 expression, and their stable MIR211-overexpressing counterparts A375/211 cells (**Supplementary Fig. S1**), were injected subcutaneously into the flanks of severe combined immunodeficiency (SCID) mice. Ectopic MIR211 expression significantly and rapidly increased aggressive tumor growth and weight (Fig. 1A-C), with rapid growth of A375/211 tumors from day six to approximately eight-times the volume of their parental controls by day 12, when mice were sacrificed due to tumor burden. An additional two independent stable clones (A375/211-c2 and A375/211-c3) behaved similarly (**Supplementary Fig. S2**), excluding a clonal effect. A negative control MIR210 (known as a hypoxia inducible miRNA and also the fine-tuner of the hypoxic response(38) did not affect tumor growth when overexpressed in A375 (A375/210) cells (**Supplementary Fig. S3A-C**). Further, to investigate whether the effect of MIR211 might be specific to A375 cells, we ectopically expressed MIR211 in another amelanotic melanoma cell line, LOX-IMV1 (LOX-IMV1/211), which also harbors a *BRAF*^V600E^ mutation(39). Again, there was a significant increase in tumor size and weight relative to the vector only control cell line *in vivo* (**Supplementary Fig. S4**).

**Figure 1.**
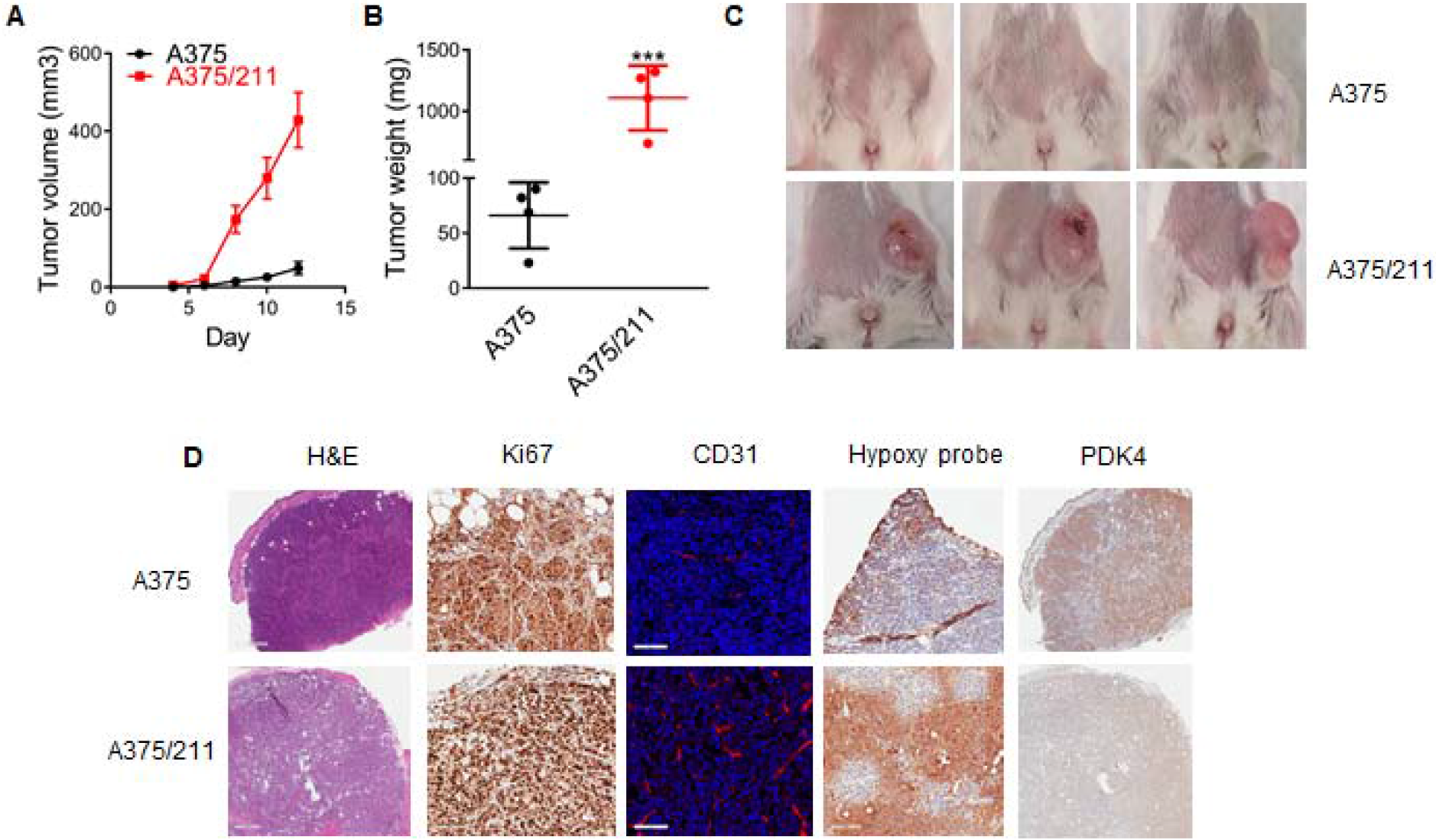
MIR211 promotes tumor growth and the cancer phenotype *in vivo*. A. A375 amelanotic melanoma cells and their MIR211 (A375/211) overexpressing counterparts were injected subcutaneously into the flanks of SCID mice and tumor volume measured with electronic calipers until termination of the experiment at 12 days. A375/211 tumors were significantly larger and grew more rapidly than the other tumors. B. Tumors were excised and weighed at the termination of the experiment on day 12. MIR211-overexpressing tumors were significantly heavier than either parental A375 xenografts. Student *t*-test, *** p ≤ 0.001. C. Photographs of subcutaneous tumor xenografts at the termination of the experiment at day 12. A375/211 xenografts were large and began to ulcerate while tumors in the other groups were barely visible. D. Photomicrographs of A375 and A375/211 xenografts showing tumor morphology (H&E; scale bar 500 um); increased proliferation in A375/211 xenografts (Ki67 IHC; scale bar 100 um); increased blood vessel density in A375/211 xenografts (CD31 IF; scale bar 100 um); increased areas of hypoxia in A375/211 xenografts (hypoxyprobe; scale bar 100 um); and decreased PDK4 expression in A375/211 xenografts (PDK4 IHC; scale bar 500 um).

Cell proliferation was examined in xenograft tissues using antibodies targeting Ki67, a pan-cell cycle marker. Approximately 30% of A375 cells showed strong nuclear Ki67 expression compared to 100% of A375/211 cells (Fig. 1D), consistent with the observation that A375/211 cells divide more rapidly than parental A375 cells. CD31 immunofluorescence (IF) to assess blood vessel density revealed that angiogenesis was markedly enhanced in A375/211 tumors compared to A375 tumors (Fig. 1D). Using pimonidazole (Hypoxyprobe ^TM^) to evaluate tumor hypoxia(40), A375/211 xenografts contained more hypoxic areas compared to A375 xenografts, suggesting that A375/211 tumors rapidly outgrew their blood supplies even with enhanced angiogenesis (Fig. 1D). We previously established a link between MIR211 and metabolism(16,41). MIR211 overexpression increases mitochondrial respiration by targeting pyruvate dehydrogenase kinase 4 (PDK4)(16), which generates mitochondrial acetyl-coA and proliferative dependence on the TCA cycle and oxidative phosphorylation(16). Consistent with these previous findings, PDK4 expression was reduced in A375/211 cells, both at the transcript (**Supplementary Fig. S5A**) and protein levels (Fig. 1D). To examine whether restoring PDK4 in A375/211 cells overcame the effects of MIR211 overexpression, stable A375/211/PDK4-re-expressing cells with similar PDK4 expression to parental A375 cells (**Supplementary Fig. S5B**) were injected into SCID mice. As expected, PDK4 activity reversed the phenotype, reducing tumor growth to below that of the parental A375 xenografts (**Supplementary Fig. S5C**). Ectopic expression of PDK4 alone in A375 cells was sufficient to rescue the aggressive phenotype in A375/211 and A375 tumors (**Supplementary Fig. S6A-C**). Interestingly, A375/PDK4 xenografts were not large until day 73 (data not shown).

### Human MIR211 in tumor xenografts modulates mouse gene expression

The oncogenic effects of MIR211 *in vivo* are paradoxical to the tumor-suppressing effects seen *in vitro*(15). To resolve this paradox, we reasoned that this may be caused by an altered tumor microenvironment as MIR211 in xenografts might either directly or indirectly interact with the host (murine) cells. This model has three essential postulates: (1) a human-specific signal is emitted by the implanted human cells; (2) mouse cells respond to the human-specific signal; (3) human cells in turn respond to mouse-specific feedback signals. Since exosomes mediate intercellular communication in tissues and contain miRNAs(42) (including MIR211(43)), we hypothesized that MIR211 might be delivered to adjacent mouse cells in the tumor microenvironment, and thereby satisfies the first condition of the model.

To test this hypothesis, A375 and A375/211 cells were grown with exosome-free fetal bovine serum, conditioned media collected, and MIR211, MIR16, and MIR23a (the latter two as positive controls(44,45)) levels analyzed by qRT-PCR. The expression levels of MIR16 and MIR23a were similar in the cellular and exosomal fractions of A375 or A375/211 cell lines (**Supplementary Fig. S7A & B)**. By contrast, as anticipated by the model, MIR211 levels were enriched approximately 100-fold in A375/211 cell-free medium (Fig. 2A) and approximately 1300-fold in A375/211 exosomes compared to the parental A375 line (Fig. 2B).

**Figure 2.**
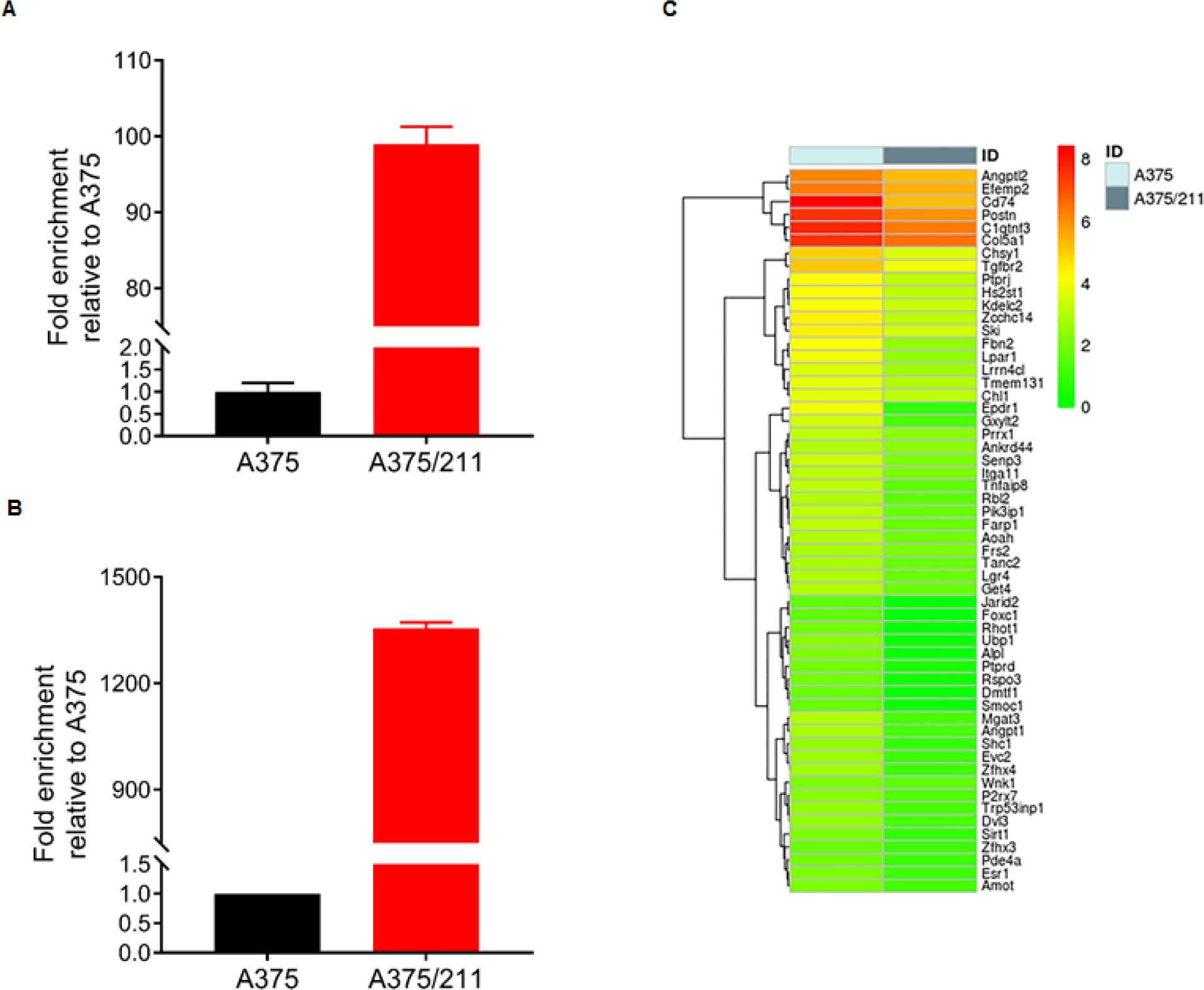
Cell-free (A) and exosome-loaded (B) MIR211 in A375/211 cells. A and B. MIR211 levels were measured in A375/211 cell free media (A) and exosomes (B) in culture and compared to parental cells. qRT-PCR results demonstrate elevated levels of MIR211 in both cell free media and exosomes. C. Heatmap for 56 putative MIR211 mouse targets showing downregulation in A375/211 xenografts.

We now examine the second postulate of the model, in which mouse cells are assumed to respond to the human cells. Since RNA-seq can accurately differentiate human and mouse transcripts in xenografts(46), we next performed RNA-seq of total RNA from xenografts to examine whether the human tumor cells affected the mouse transcriptome. First, reads were separately aligned to the human (GRCH37/hg19) and mouse (GRCm/mm10) genomes, after which only mouse transcriptome changes were assessed by Ingenuity Pathway Analysis (IPA) to identify the effect of human cells on the mouse transcriptome. IPA analysis revealed a number of activated pathways including cell cycle, growth, proliferation, movement, and cardiovascular system development and function (**Supplementary Table S1**). Next, the same sequencing data were subjected to Gene Set Enrichment Analysis (GSEA), which revealed enrichment of gene expression related to hypoxia response, angiogenesis and endothelial cell pathways in the mouse transcriptome **(Supplementary Fig. S8**), consistent with the increased CD31 endothelial staining seen in A375/211 xenografts between cell junctions (Fig. 1D).

We next analyzed the expression levels of putative MIR211 targets in mouse RNA-seq data. Human and mouse MIR211 sequences differ by only a single nucleotide towards the 3’ end (**Supplementary Fig. S7C**), so their targets are likely to be similar; indeed, using an miRNA target prediction algorithm(37), human and mouse MIR211 had the same predicted targets (data not shown). Among 481 putative targets common to both target prediction algorithms (microRNA.org(36) and miRDB(37)), 56 mouse genes were significantly downregulated (Fig. 2C **and Supplementary Table S2**, log_2_ fold change ≤ −1.0, p < 0.05) including *Angpt1*, which is often decreased in cancer and associated with increased vasculogenesis in renal cell carcinoma(47). Similar to our results, *Dror et al.* recently reported that MIR211 is transferred to fibroblasts through melanosomes to promote a cancer-associated fibroblast phenotype(11). Taken together, these results support the possibility that MIR211 is transferred to adjacent mouse endothelial cells via exosomes to participate in altering the cellular properties of mouse cells to those previously associated with various cancers.

### Transcriptomic changes in xenografted human cells within the tumor microenvironment

Explicit testing of the feedback effects of murine cells on the human tumor cells is difficult in the absence of an identified feedback effector molecule. We therefore used an indirect measure and investigated whether MIR211 expression in xenografts causes changes in the human-specific transcriptome relative to that in the culture tumor cells that are sufficient to explain rapid tumor growth phenotype.

A375, A375/211, and A375/211/PDK4 xenografts were subjected to RNA-seq and human-specific transcriptomes were analyzed (**Supplementary Table S3 and Supplementary Fig. S9**). The results were consistent with the activation of four human-specific pathways (cyclins and cell cycle regulation, ERK5 signaling, hypoxia signaling, and telomerase signaling; z-score ≥ 2.0) and the inactivation of five human-specific pathways (cAMP-mediated signaling, interferon signaling, PPARα/RXRα activation, Gαi signaling, and Gαq signaling; z-score ≤ −2.0) in tumors expressing the human-specific MIR211. These results were not previously observed in A375 cells or A375/211 cells grown *in vitro* cultures (30).

To determine whether upregulated genes in different xenografts (log_2_ fold change ≥ 1.0, p ≤ 0.05) shared common biological functions, we performed GSEA(48) on the human-specific upregulated transcripts. Epithelial-to-mesenchymal transition (EMT) was the most significantly enriched gene set (**Supplementary Table S4**, FDR = 4.87 x 10^−27^). In addition, MIR211 altered several EMT-related pathways including estrogen response genes (early (FDR = 1.52 x 10^−2^) and late (FDR = 8.33 x 10^−4^)), genes upregulated in hypoxia (FDR = 4.09E x 10^−23^), and genes responding to ultraviolet (UV) radiation (up (FDR = 2.15E x 10^−4^) and down (FDR = 2.41 x 10^− 6^))(49,50). Several lipid metabolism-related gene sets including cholesterol homeostasis, adipogenesis, and fatty acid metabolism were also upregulated (FDR = 2.54 x 10^−6^, 1.63 x 10^−4^, and 2.15 x 10^−4^, respectively).

### MIR211 directly targets negative regulators of the ERK5 pathway

Four human-specific signaling pathways were activated in A375/211 compared to A375 xenografts (**Supplementary Table S3 and Supplementary Fig. S9**) including the ERK5 signaling axis. ERK5 itself was recently shown to be activated in melanomas(51), and we reasoned that negative regulators of the ERK5 pathway could be MIR211 targets. Using TargetScan(52), microRNA.org(36), and miRDB(37), we identified putative MIR211 targets in the ERK5 signaling pathway, which included baculoviral IAP repeat containing 2 (*BIRC2*) and several dual specificity phosphatases (DUSPs) including *DUSP3, DUSP4, DUSP6*, and *DUSP19*. BIRC2 is an ubiquitin ligase and its family member, BIRC4, is a known regulator of MEKK2, which lies upstream of MAPK kinase kinase in the ERK5 pathway(53). DUSP proteins are possible phosphatases of ERK5 signaling proteins, which are expected to inhibit ERK5 signaling activation (**Supplementary Fig. S10**), and DUSP4 was previously predicted as an MIR211 target gene in melanoma(54).

In an attempt to determine which candidates were directly regulated by MIR211 *in vivo*, we next directly determined human-specific MIR211 targets using Argonaute-2 (Ago2) immunopurification, with co-immunopurified RNAs determined by RNA-seq. To validate the RIP-seq results, bound RNAs were filtered by MIR211-binding sites to identify putative MIR211 targets. Both *BIRC2* and *DUSP* family members were enriched by Ago2 immunopurification (**Supplementary Table S5**).

We next performed an miRNA target cleavage assay (luciferase reporter assay) to further confirm that DUSP6 (Fig. 3A) and BIRC2, DUSP4, and DUSP19 are direct MIR211 targets (**Supplementary Fig. S11A**). When reporter plasmids were co-transfected with plasmids expressing MIR211 into HEK-293T cells, the wild-type DUSP6 reporter showed reduced luciferase activity, while the mutant one did not (Fig. 3A). Further, when a reporter carrying luciferase with the 3’UTR of DUSP6 was transiently transfected to A375 or A375/211 cells, a reduction in luciferase activity was observed in A375/211 cells compared to A375 cells (Fig. 3B). To determine which DUSPs specifically dephosphorylate ERK5, A375 cells were treated with siRNAs against DUSP3, DUSP4, DUSP6, and DUSP19, and ERK5 phosphorylation was assessed. As shown in **Supplementary Fig. S11B**, knockdown of each DUSP except DUSP3 increased ERK5 phosphorylation. Furthermore, when DUSP proteins were ectopically expressed in A375/211 cells, DUSP6 rapidly decreased ERK5 phosphorylation, indicating DUSP6 may act rapidly and specifically on ERK5 (Fig. 3C **and Supplementary Fig. S11C**). DUSP6 protein level was lower in A375/211 cells and corresponded to higher ERK5 phosphorylation than in A375 cells (Fig. 3D). Next we engineered a cell line to re-express DUSP6 in A375/211 background and cells were injected subcutaneously into the flanks of SCID mice together with cells from A375 and A375/211 as controls. The re-expression of DUSP6 significantly reduced A375/211 tumor growth (Fig. 3E), and DUSP6-overexpressing tumors were significantly smaller compared to A375/211 xenografts harvested the same day (Fig. 3F; p ≤ 0.001). Given these results, we further investigated the role of DUSP6 and ERK5 axes in melanoma.

**Figure 3.**
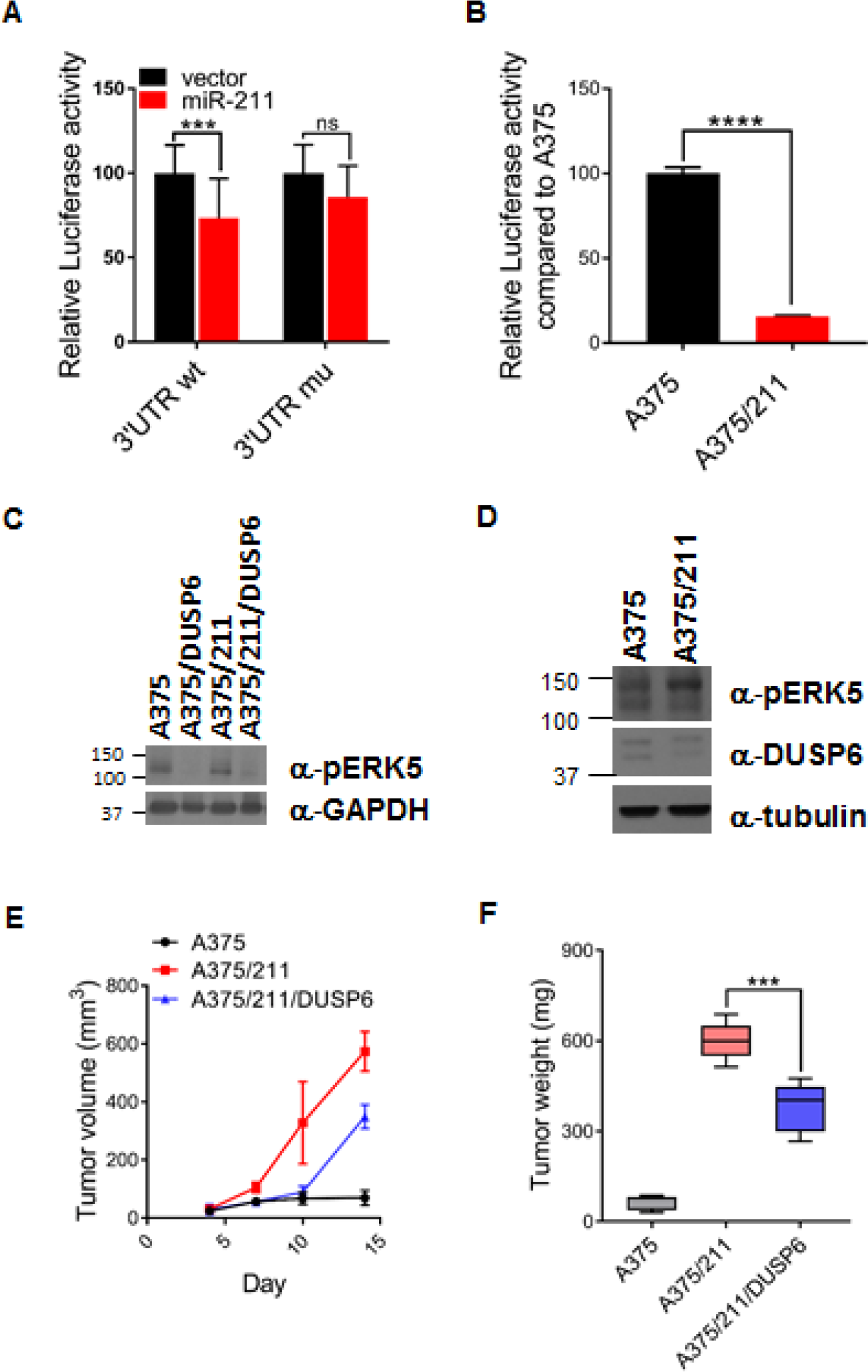
MIR211 targets negative regulators of ERK5 pathway. A. Luciferase reporter assays carrying either wild-type 3’UTR or a mutant with several nucleotides swapped in the region corresponding to the MIR211 seed. When reporter plasmids were co-transfected with plasmids expressing MIR211 into HEK-293T cells, the wild-type DUSP6 reporter showed reduced luciferase activity, while the mutant one did not. Student *t*-test, ns: not significant, *** p ≤ 0.001. B. Reporter carrying luciferase with 3’UTR of DUSP6 was transiently transfected to A375 or A375/211 cells. A reduction in luciferase activity was observed in A375/211 cells compared to A375 cells. Student *t*-test, **** p ≤ 0.0001. C. Western blot showing the decrease of phospho-ERK5 (pERK5) upon overexpression of DUSP6. D. Western blot analysis of pERK5 and DUSP6 in A375 and A375/211 cells E. A375, A375/211, and A375/211/DUSP6 cells were injected subcutaneously into the flanks of SCID mice and tumor volume measured with electronic calipers until termination of the experiment at 14 days. The ectopic expression of DUSP6 reduced significantly A375/211 tumor growth. F. Tumors were excised and weighed at the termination of the experiment at 14 days. DUSP6-overexpressing tumors were significantly lighter than A375/211 xenografts. Student t-test, *** p ≤ 0.001.

### ERK5 pathway activation by MIR211 confers melanoma drug resistance

To determine if DUSP6 affects ERK1/2, we analyzed ERK1/2 phosphorylation upon DUSP6 siRNA knockdown (Fig. 4A**, Supplementary Fig. S12**); ERK1/2 phosphorylation was not affected by DUSP6 knockdown (Fig. 4A).

**Figure 4.**
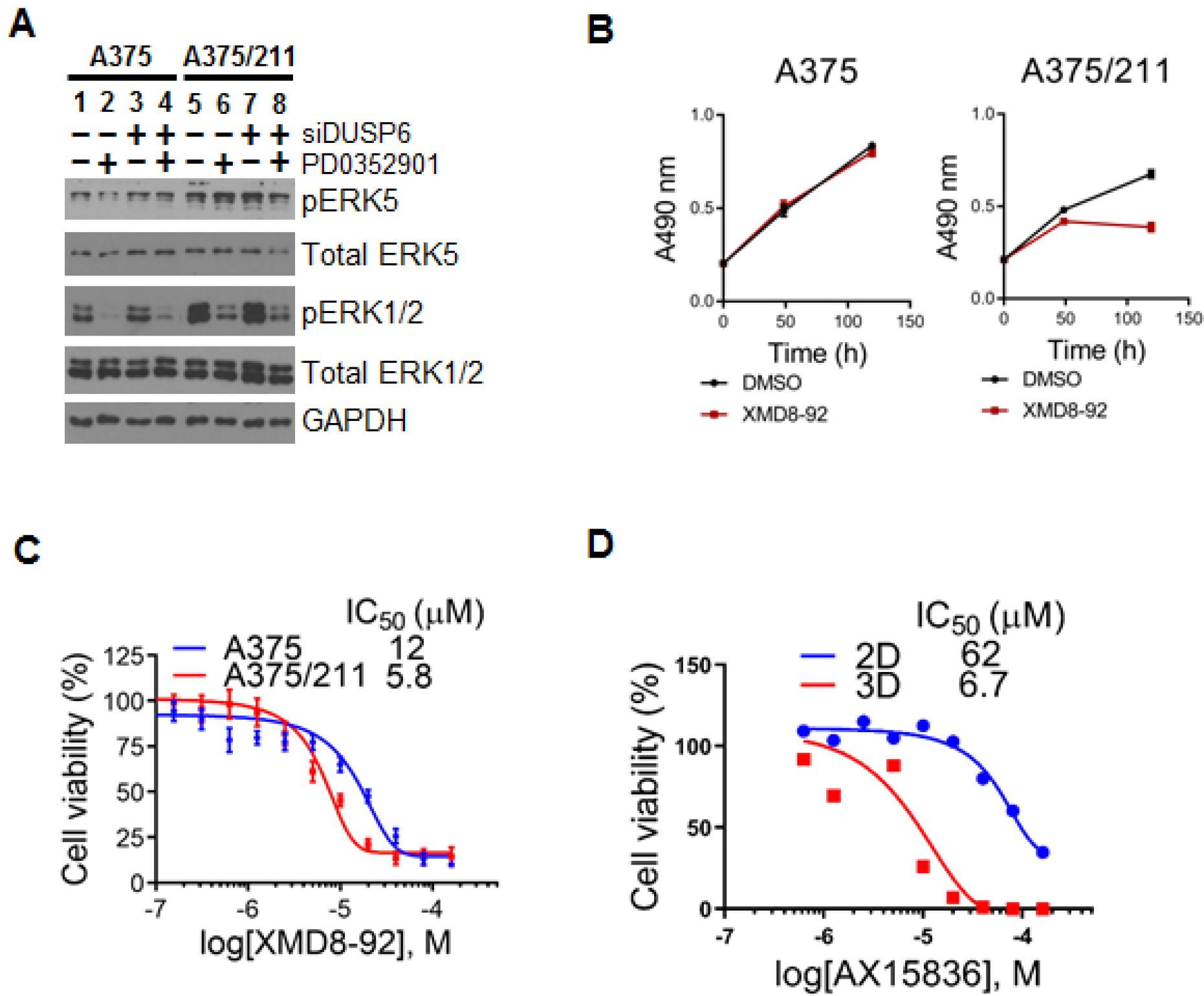
MIR211 activates the ERK5 pathway via DUSP6. A. Western blots of A375 and A375/211 cells treated with siDUSP6 or PD0325901 (100 nM), a MEK1/2 inhibitor. Phospho-ERK5 (pERK5) was lower in A375 cells than A375/211 cells and increased upon siDUSP6 treatment in both cell types. ERK1/2 phosphorylation was not affected by *DUSP6* knockdown. PD0325901 efficiently abrogated ERK1/2 phosphorylation and slightly increased ERK5 phosphorylation. B. Cell viability assays showing that A375/211 cells were more sensitive to the ERK5 inhibitor XMD8-92, suggesting that A375/211 cells are more dependent on the ERK5 pathway. C. IC50 concentration of XMD8-92 for A375 and A375/211 cells. XMD8-92 IC50 value of A375/211 cells was about two folds lower than that of A375. D. IC50 concentration of AX15836 for A375/211 cells in two and three dimensional culture. In 3D culture, A375/211 cells were much sensitive than 2D culture to AX15836, while A375 and A375/211 cells showed similar sensitivity to AX15836 in 2D culture.

Phospho-ERK5 (pERK5) was lower in A375 cells than in A375/211 cells and increased upon siDUSP6 treatment in both cell lines (Fig. 4A). As expected, stable overexpression of DUSP6 decreased phospho-ERK5 levels (Fig. 3C). ERK5 is activated upon ERK1/2 inhibition in intestinal epithelial cells and colorectal cancer cell lines(55). To establish whether ERK5 is also activated by ERK1/2 pathway inhibition in melanoma, A375 and A375/211 cells were treated with PD0325901, which inhibits the ERK1/2 kinase MEK1/2. PD0325901 efficiently abrogated ERK1/2 phosphorylation and slightly increased ERK5 phosphorylation (Fig. 4A). Since the ERK5 pathway was activated in A375/211 cells due to DUSP6 inhibition by MIR211, we hypothesized that the ERK5 pathway may compensate for loss of ERK1/2 signaling to drive hyperproliferation in melanomas. Consistent with this, A375/211 cells were more sensitive to the ERK5 inhibitor XMD8-92 than parental cells (Fig. 4B & 4C). XMD8-82, the first generation ERK5 inhibitor, has been reported to have off-target activity on bromodomains(56), so we examined the effect of AX15836, an XMD8-92 derivative with no significant off-target activity(56) on A375/211 cell proliferation. While AX15836 had no anti-proliferative effect on two-dimensional cultures over a short period (48 h), when cells were grown in 3-D cultures (colony forming assay), the IC50 of AX15836 on A375/211 cells was reduced from 62 uM to 6.7 uM, indicating that A375/211 cells were more sensitive to drug than A375 cells (Fig. 4D). Further, A375/211 cell growth was inhibited by ERK5 siRNA knockdown (**Supplementary Fig. S13B**).

Finally, MIR211 expression increases in melanomas treated with the BRAF inhibitor vemurafenib and may contribute to vemurafenib resistance(27,28,43). MIR211 levels were similarly increased in A375 cells treated with vemurafenib (Fig. 5A) and the MEK inhibitor cobimetinib (Fig. 5B). In addition, ectopic MIR211 expression increased vemurafenib resistance 10.4-fold compared to A375 cells (Fig. 5C). To examine whether vemurafenib resistance may be mediated via the MIR211-ERK5 axis, pERK5 levels were evaluated in vemurafenib-resistant cell lines (Fig. 5D), previously thought to become resistant primarily due to ERK1/2 pathway reactivation(27). Indeed, while ERK1/2 phosphorylation was significantly increased in resistant cell lines compared to parental A375 cells, ERK5 phosphorylation also increased, suggesting that ERK5 signaling might also contribute to acquired drug resistance in melanoma. In agreement with this hypothesis, MIR211 expression was also significantly increased in these cell lines (Fig. 5E). To further investigate these observations, MIR211 gene expression was quantified in 39 melanoma cell lines (**Supplementary Table S6**). Interestingly, MIR211 expression (low and high) and vemurafenib resistance partitioned melanoma cell lines into two groups (**Supplementary Fig. S14**). Vemurafenib resistance and MIR211 expression were correlated in the MIR211 low but not the MIR211-high group (Fig. 5F).

**Figure 5.**
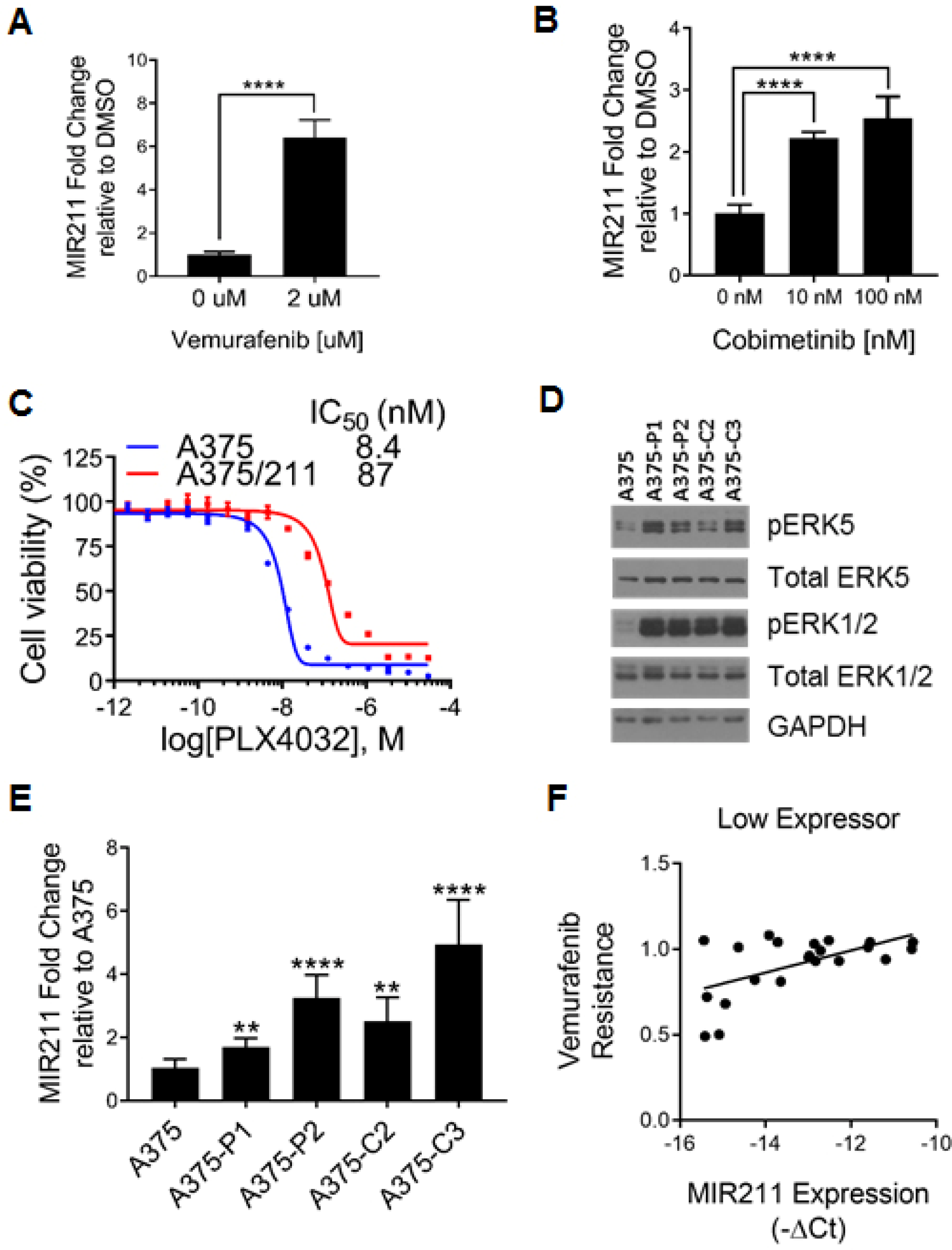
MIR211 contributes to vemurafenib resistance via ERK5. A. MIR211 expression levels increased in A375 cells treated with vemurafenib. B. MIR211 levels increased in A375 cells treated with MEK1 inhibitor, cobimetinib. C. Cell viability assays showing that ectopic MIR211 expression increased vemurafenib resistance by 10.4-fold compared to A375 cells. D. Both ERK1/2 and ERK5 phosphorylation increase markedly and significantly in four clones of A375 cells with acquired vemurafenib resistance. E. MIR211 levels increased in vemurafenib resistant cell lines compared to A375 cells. F. Vermurafenib resistance and MIR211 levels were correlated in melanoma cell lines with low expression level of MIR211. (Pearson r = 0.5781, 95% confidence interval = 0.207 to 0.8038, R^2^ = 0.3343, p = 0.0048).

### *In vivo* tumor growth induced by MIR211 is not fully explained by metabolic changes

PDK inhibits pyruvate dehydrogenase(57) and, in cancer, PDK shunts glucose from TCA cycle to produce lactate(58). We previously showed that inhibition of PDK4 by MIR211 overexpression in A375 cells *in vitro* increased the oxygen consumption rate (OCR), indicating enhanced mitochondrial activity(16). To determine whether PDK4 regulation affects tumor growth by modulating metabolism and whether this effect persists *in vivo*, we performed targeted metabolite analysis (**Supplementary Fig. S15A**). Metabolites were extracted from A375, A375/211, or A375/211/PDK xenografts, derivatized (described above), and analyzed by mass spectrometry.

MIR211 overexpression did not significantly affect glycolysis intermediates, although ectopic PDK4 expression significantly decreased lactate production (**Supplementary Fig. S15B**). Similarly, TCA cycle intermediates were generally unaffected except for succinate (reduced by PDK4-re-expression) and fumarate (decreased by MIR211 overexpression but partially restored in A374/211/PDK4 xenografts) (**Supplementary Fig. S15C**). Amino acid levels were almost uniformly decreased in A375/211 xenografts and partially restored in A375/211/PDK4 xenografts except for threonine, which further decreased with the addition of PDK4 (**Supplementary Fig. S15D**). Overall, although MIR211 acts as a metabolic switch *in vitro*(16) and in that context might dictate the cellular phenotype, MIR211’s effects on metabolic pathways appear to be largely “buffered” within the physiological environment *in vivo*. It is still possible that human-specific exosomal MIR211 does switch murine cell metabolism (Warburg effects), seen previously in cultured cells(30), and the feedback effects by the mouse cells prevent this metabolic switch to be actuated in the human cells.

In conclusion, these results suggest that MIR211 has pleiotropic effects *in vitro* and *in vivo*.

While MIR211 appears to render melanoma cells metabolically vulnerable *in vitro*, these effects appear to be largely buffered *in vivo*, instead upregulating critical cancer pathways including ERK5 signaling to regulate the cell cycle and angiogenesis, with therapeutic consequences. Our results are summarized in the model we propose in Figure 6. As we proposed earlier, a human-specific signal is released with MIR211 from the implanted human cells via exosome to mouse cells in turn respond to mouse-specific feedback signals. Melanoma cells appear to adapt their microenvironment and adapt to their microenvironment to gain a growth advantage, especially when drugs apply selection pressure. It remains to be determined what factors dictate which pathways are targeted by miRNAs at any given point in time, but future precision therapies based on MIR211 or its target pathways are likely to require consideration of this exquisite context-dependency.

**Figure 6.**
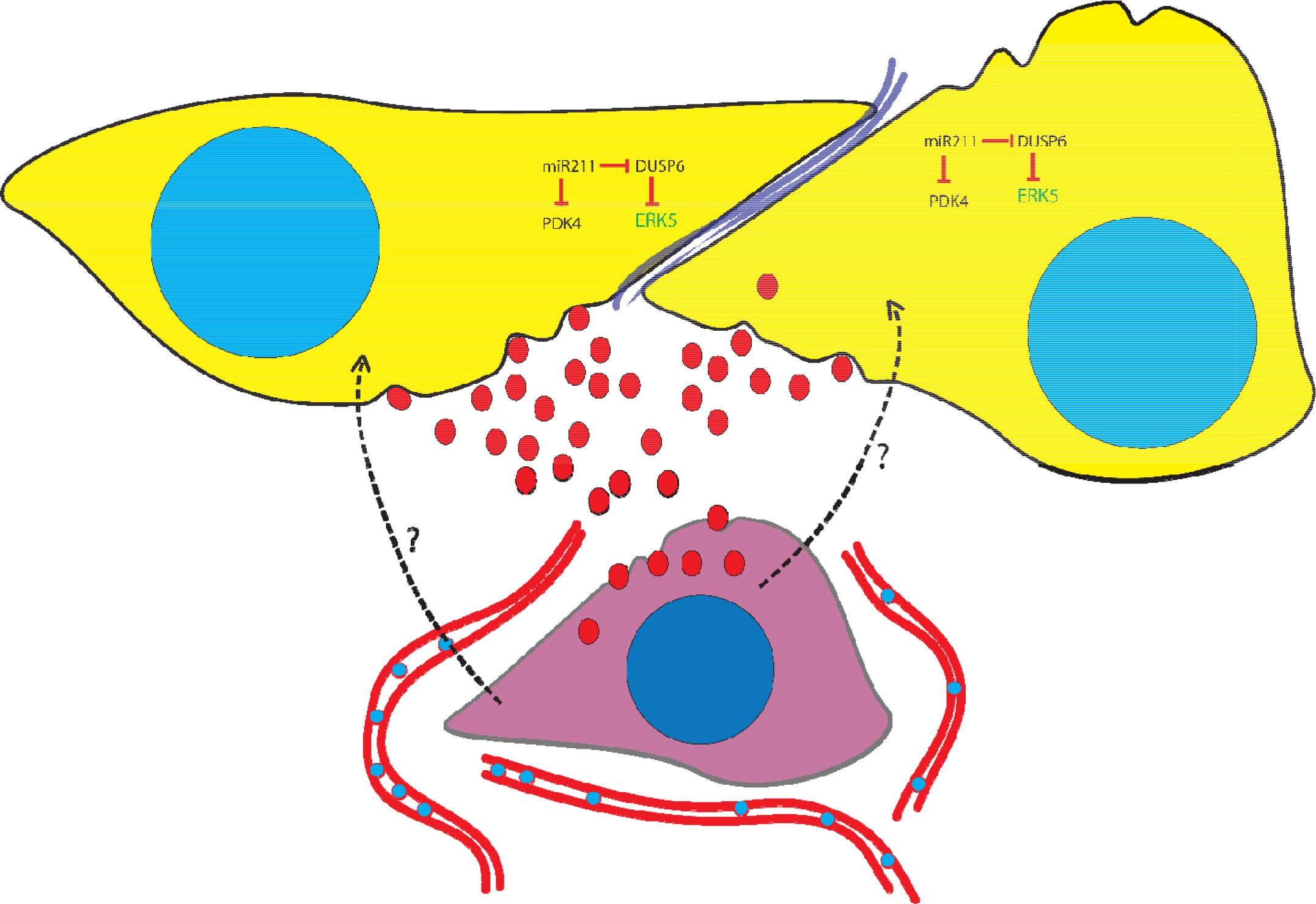
A model summarizing the interaction between A375/211 and the tumor microenvironment. Human tumor cells (yellow) expressing MIR211, secrete exosomes (red ellipses) loaded with MIR211, which enter mouse cells (purple). In mouse cells, MIR211 inhibits its murine targets, which in turn activate the transcription of a number genes including ANGPTL2, EFEMP2, CD74, POSTN, C1QTNF3, and COL5A1. These changes in murine cells stimulate angiogenesis and cause feedback to the human cells, interacting with the MIR211 circuit involving DUSP6, to induce ERK5 signaling, resulting in tumor hyper-proliferation. While MIR211’s target PDK4 is inhibited within human cells, which can be rescued by over-expressing the *PDK4* cDNA.

## ACCESSION NUMBERS

All xenografts RNA-seq and Ago2 RIP-seq datasets have been deposited with Gene Expression Omnibus (GEO) under accession number GSE125836.

## ACKNOWLEDGEMENTS

We thank Sanford Burnham Prebys Medical Discovery Institute Analytical Genomics core, Bioinformatics core, Histology, and Microscope core and Metabolomics core for their support. We also thank Drs. Jeffrey M. Trent (Translational Genomics Institute, Phoenix, AZ) originally provided the 33 UACC cell lines and Neal Rosen (Memorial Sloan Kettering Cancer Center, New York, NY) provided the 5 SK-MEL and MeWo cell lines for the studies.

## Footnotes

a) Ding, K.-F., Finlay, D., Yin, H., Hendricks, W.P.D., Sereduk, C., Kiefer, J., Sekulic, A., LoRusso, P.M., Vuori, K., Trent, J.M. and Schork, N.J. (2017) Analysis of Variability in High Throughput Screening Data Applications to Melanoma Cell Lines and Drug Responses. *Oncotarget* 8:27786-27799. doi: 10.18632/oncotarget.15347

b) Ding, K.-F., Petricoin III, E., Finlay, D., Yin, H., Hendricks, W.P.D., Sereduk, C., Kiefer, J., Sekulic, A., LoRusso, P.M., Vuori, K., Trent, J.M. and Schork, N.J. (2017) Nonlinear Mixed Effects Dose Response Modeling in High Throughput Drug Screens: Application to Melanoma Cell Analysis. *Oncotarget* 9:5044-5057. doi: 10.18632/oncotarget.23495

c) Ding, K.-F., Finlay, D., Yin, H., Hendricks, W.P.D., Sereduk, C., Kiefer, J., Sekulic, A., LoRusso, P.M., Vuori, K., Trent, J.M. and Schork, N.J. (2018) Network Rewiring in Cancer: Applications to Melanoma Cell Lines and the Cancer Genome Atlas Patients. *Front. Genet*. 9:228. doi: 10.3389/fgene.2018.00228

