## Supplementary file for "Exosome-mediated MIR211 modulates tumor microenvironment via the DUSP6-ERK5 axis and contributes to BRAFV600E inhibitor resistance in melanoma"

| Categories | Diseases or Functions Annotation | p-value | Predicted Activation State | Activation z-score |
|---|---|---|---|---|
| Cell Cycle | M phase of tumor cell lines | 0.0000 | Increased | 2.712 |
| | M phase | 0.0000 | Increased | 2.591 |
| | M phase of cervical cancer cell lines | 0.0000 | Increased | 2.126 |
| Immunological Disease | systemic autoimmune syndrome | 0.0000 | Increased | 2.362 |
| Cardiovascular System Development and Function | development of vasculature | 0.0000 | Increased | 2.239 |
| | angiogenesis | 0.0000 | Increased | 2.239 |
| | development of vascular endothelial cells | 0.0000 | Increased | 2.226 |
| | cell movement of endothelial cells | 0.0000 | Increased | 2.171 |
| Cardiovascular Disease | vascular lesion | 0.0000 | Increased | 2.000 |
| Connective Tissue Disorders, Inflammatory Disease, Organismal Injury and Abnormalities, Skeletal and Muscular Disorders | Rheumatic Disease | 0.0000 | Increased | 2.152 |
| Cellular Development, Cellular Growth and Proliferation | cell proliferation of carcinoma cell lines | 0.0000 | Increased | 2.101 |
| Cellular Movement | cell movement of tumor cell lines | 0.0000 | Increased | 2.087 |

The activation z-score measures the match between expected relationship direction and observed gene expression (≥2 is considered significant).

**Supplementary Table S1.** Ingenuity Pathway Analysis (IPA) of mouse gene expression in MIR211-overexpressign xenografts compared to A375 parental xenografts.

| Gene | A375/211 (Reads) | A375 (Reads) | A375/211 (RPKM) | A375 (RPKM) | logFC ( 211/A375) | p-value | FDR |
|---|---|---|---|---|---|---|---|
| Alpl | 0.6 | 43.5 | 0.2 | 3.2 | -3.7 | 0.0007 | 0.01 |
| Amot | 15.0 | 124.0 | 1.4 | 3.3 | -1.4 | 0.0179 | 0.14 |
| Angpt1 | 9.3 | 129.2 | 1.4 | 5.6 | -2.2 | 0.0006 | 0.01 |
| Angptl2 | 201.0 | 1407.0 | 38.1 | 76.0 | -1.2 | 0.0118 | 0.10 |
| Ankrd44 | 36.0 | 239.0 | 3.8 | 7.2 | -1.1 | 0.0362 | 0.22 |
| Aoah | 13.1 | 103.7 | 2.9 | 6.6 | -1.4 | 0.0293 | 0.19 |
| C1qtnf3 | 277.4 | 2776.8 | 77.0 | 219.9 | -1.7 | 0.0003 | 0.01 |
| Cd74 | 83.8 | 2701.3 | 38.6 | 355.3 | -3.4 | 0.0000 | 0.00 |
| Chl1 | 87.0 | 560.0 | 7.3 | 13.4 | -1.1 | 0.0284 | 0.19 |
| Chsy1 | 78.0 | 715.0 | 12.2 | 31.9 | -1.6 | 0.0014 | 0.02 |
| Col5a1 | 1136.0 | 7956.0 | 88.2 | 176.3 | -1.2 | 0.0100 | 0.09 |
| Dmtf1 | 1.3 | 50.4 | 0.2 | 2.5 | -4.0 | 0.0006 | 0.01 |
| Dvl3 | 7.0 | 70.0 | 1.5 | 4.4 | -1.7 | 0.0189 | 0.14 |
| Efemp2 | 107.9 | 666.5 | 45.2 | 79.7 | -1.0 | 0.0357 | 0.22 |
| Epdr1 | 3.0 | 181.0 | 0.8 | 14.1 | -4.3 | 0.0000 | 0.00 |
| Esr1 | 11.5 | 117.7 | 1.2 | 3.5 | -1.7 | 0.0071 | 0.07 |
| Evc2 | 7.0 | 119.0 | 1.1 | 5.4 | -2.5 | 0.0003 | 0.01 |
| Farp1 | 18.0 | 194.0 | 2.4 | 7.4 | -1.8 | 0.0016 | 0.02 |
| Fbn2 | 77.0 | 922.0 | 4.8 | 16.4 | -2.0 | 0.0001 | 0.00 |
| Foxc1 | 0.0 | 41.0 | 0.0 | 1.9 | -7.8 | 0.0001 | 0.00 |
| Frs2 | 28.0 | 189.0 | 3.2 | 6.2 | -1.2 | 0.0372 | 0.22 |
| Get4 | 7.5 | 72.2 | 2.3 | 6.4 | -1.6 | 0.0315 | 0.20 |
| Gxylt2 | 4.0 | 99.0 | 1.4 | 10.1 | -3.0 | 0.0001 | 0.00 |
| Hs2st1 | 59.0 | 457.0 | 8.0 | 17.7 | -1.3 | 0.0075 | 0.07 |
| Itga11 | 25.0 | 203.0 | 3.3 | 7.7 | -1.4 | 0.0110 | 0.09 |
| Jarid2 | 0.4 | 68.1 | 0.0 | 2.2 | -8.5 | 0.0000 | 0.00 |
| Kdelc2 | 48.0 | 345.0 | 8.6 | 17.6 | -1.2 | 0.0165 | 0.13 |
| Lgr4 | 18.0 | 182.0 | 2.3 | 6.7 | -1.7 | 0.0029 | 0.03 |
| Lpar1 | 23.9 | 267.5 | 4.6 | 14.8 | -1.9 | 0.0007 | 0.01 |
| Lrrn4cl | 21.0 | 152.0 | 5.3 | 11.0 | -1.3 | 0.0329 | 0.21 |
| Mgat3 | 10.0 | 160.0 | 1.4 | 6.4 | -2.4 | 0.0001 | 0.00 |
| P2rx7 | 6.4 | 56.5 | 1.6 | 4.1 | -1.6 | 0.0361 | 0.22 |
| Pde4a | 8.3 | 72.6 | 1.2 | 2.9 | -1.6 | 0.0271 | 0.18 |
| Pik3ip1 | 8.0 | 94.0 | 2.3 | 7.8 | -1.9 | 0.0045 | 0.05 |
| Postn | 314.0 | 3045.3 | 63.7 | 176.4 | -1.7 | 0.0005 | 0.01 |
| Prrx1 | 30.7 | 213.7 | 4.0 | 8.0 | -1.2 | 0.0292 | 0.19 |
| Ptprd | 7.0 | 139.0 | 0.5 | 2.8 | -2.7 | 0.0001 | 0.00 |
| Ptprj | 93.4 | 655.6 | 8.0 | 16.0 | -1.2 | 0.0139 | 0.11 |
| Rbl2 | 16.5 | 189.6 | 2.2 | 7.2 | -1.9 | 0.0012 | 0.02 |
| Rhot1 | 0.0 | 65.6 | 0.0 | 3.0 | -8.5 | 0.0000 | 0.00 |
| Rspo3 | 1.0 | 35.0 | 0.3 | 2.7 | -3.4 | 0.0034 | 0.04 |
| Senp3 | 12.0 | 114.2 | 3.2 | 8.7 | -1.6 | 0.0090 | 0.08 |
| Shc1 | 4.5 | 81.7 | 0.9 | 4.8 | -2.4 | 0.0013 | 0.02 |
| Sirt1 | 4.7 | 75.7 | 0.8 | 3.6 | -2.3 | 0.0026 | 0.03 |
| Ski | 85.0 | 627.0 | 10.1 | 21.4 | -1.3 | 0.0100 | 0.09 |
| Smoc1 | 1.2 | 44.3 | 0.2 | 2.4 | -3.8 | 0.0007 | 0.01 |
| Tanc2 | 47.0 | 377.0 | 2.6 | 5.9 | -1.4 | 0.0071 | 0.07 |
| Tgfbr2 | 110.3 | 921.3 | 15.0 | 35.8 | -1.5 | 0.0029 | 0.03 |
| Tmem131 | 66.0 | 450.0 | 6.6 | 12.8 | -1.2 | 0.0199 | 0.15 |
| Tnfaip8 | 9.7 | 142.7 | 1.9 | 7.9 | -2.2 | 0.0005 | 0.01 |
| Trp53inp1 | 12.8 | 133.9 | 1.5 | 4.6 | -1.8 | 0.0046 | 0.05 |
| Ubp1 | 1.4 | 77.2 | 0.2 | 3.7 | -4.6 | 0.0000 | 0.00 |
| Wnk1 | 30.0 | 199.6 | 1.9 | 3.5 | -1.1 | 0.0370 | 0.22 |
| Zcchc14 | 78.0 | 729.0 | 7.9 | 21.0 | -1.6 | 0.0012 | 0.02 |
| Zfhx3 | 27.0 | 214.0 | 1.1 | 2.4 | -1.4 | 0.0116 | 0.10 |
| Zfhx4 | 24.0 | 398.0 | 1.1 | 5.3 | -2.4 | 0.0000 | 0.00 |

**Supplementary Table S2.** Significantly downregulated genes in A375/211 xenografts predicted to be MIR211 targets in the mouse.

| A375/211 vs. A375 | | A375/211 vs. A375/211/PDK4 | | A375/211/PDK4 vs. A375 | |
|---|---|---|---|---|---|
| **Ingenuity Canonical Pathways** | **Z-score** | **Ingenuity Canonical Pathways** | **Z-score** | **Ingenuity Canonical Pathways** | **Z-score** |
| Antioxidant Action of Vitamin C | nd | Antioxidant Action of Vitamin C | nd | Antioxidant Action of Vitamin C | 2.000 |
| BMP signaling pathway | nd | BMP signaling pathway | nd | BMP signaling pathway | -2.236 |
| cAMP-mediated signaling | -2.000 | cAMP-mediated signaling | -2.111 | cAMP-mediated signaling | -1.000 |
| Cardiac Hypertrophy Signaling | -0.333 | Cardiac Hypertrophy Signaling | -2.309 | Cardiac Hypertrophy Signaling | 0.000 |
| Colorectal Cancer Metastasis Signaling | 0.218 | Colorectal Cancer Metastasis Signaling | -2.132 | Colorectal Cancer Metastasis Signaling | 0.000 |
| Corticotropin Releasing Hormone Signaling | 0.000 | Corticotropin Releasing Hormone Signaling | -2.000 | Corticotropin Releasing Hormone Signaling | nd |
| CREB Signaling in Neurons | 0.707 | CREB Signaling in Neurons | -2.121 | CREB Signaling in Neurons | -0.707 |
| Cyclins and Cell Cycle Regulation | 2.000 | Cyclins and Cell Cycle Regulation | nd | Cyclins and Cell Cycle Regulation | nd |
| Dopamine-DARPP32 Feedback in cAMP Signaling | -1.508 | Dopamine-DARPP32 Feedback in cAMP Signaling | -2.236 | Dopamine-DARPP32 Feedback in cAMP Signaling | -1.134 |
| EIF2 Signaling | -0.816 | EIF2 Signaling | -2.333 | EIF2 Signaling | 0.333 |
| eNOS Signaling | -1.732 | eNOS Signaling | -2.138 | eNOS Signaling | -0.302 |
| ERK5 Signaling | 2.236 | ERK5 Signaling | 0.000 | ERK5 Signaling | nd |
| Gαi Signaling | -2.121 | Gαi Signaling | -1.633 | Gαi Signaling | nd |
| Gαq Signaling | -2.333 | Gαq Signaling | -2.309 | Gαq Signaling | -0.577 |
| Hypoxia Signaling in the Cardiovascular System | 2.000 | Hypoxia Signaling in the Cardiovascular System | nd | Hypoxia Signaling in the Cardiovascular System | nd |
| Interferon Signaling | -2.449 | Interferon Signaling | -2.449 | Interferon Signaling | nd |
| P2Y Purigenic Receptor Signaling Pathway | 0.000 | P2Y Purigenic Receptor Signaling Pathway | -2.333 | P2Y Purigenic Receptor Signaling Pathway | 0.000 |
| PPARα/RXRα Activation | -2.121 | PPARα/RXRα Activation | -1.265 | PPARα/RXRα Activation | 1.342 |
| Relaxin Signaling | -0.816 | Relaxin Signaling | -2.121 | Relaxin Signaling | -0.378 |
| RhoGDI Signaling | 1.000 | RhoGDI Signaling | 1.508 | RhoGDI Signaling | -2.714 |
| Role of NFAT in Regulation of the Immune Response | -1.000 | Role of NFAT in Regulation of the Immune Response | -2.111 | Role of NFAT in Regulation of the Immune Response | -0.632 |
| SAPK/JNK Signaling | 0.000 | SAPK/JNK Signaling | -2.449 | SAPK/JNK Signaling | 0.447 |
| Sperm Motility | -0.378 | Sperm Motility | 0.000 | Sperm Motility | -2.333 |
| Synaptic Long-Term Depression | -0.707 | Synaptic Long-Term Depression | nd | Synaptic Long-Term Depression | -2.646 |
| Telomerase Signaling | 2.121 | Telomerase Signaling | 0.000 | Telomerase Signaling | 0.000 |

**Supplementary Table S3.** Ingenuity pathway analysis (IPA) of differential gene expression in A375 xenografts, MIR211-overexpressing xenografts, or MIR211/PDK4 co-expressing xenografts. Turquoise = activated; brown = inactivated. The activation z-score measures the match between expected relationship direction and observed gene expression (≥2 is considered significant).

| Gene Set Name | k/K | *p*-value | FDR |
|---|---|---|---|
| EPITHELIAL_MESENCHYMAL_TRANSITION | 0.14 | 9.74E-29 | 4.87E-27 |
| HYPOXIA | 0.13 | 1.64E-24 | 4.09E-23 |
| MTORC1_SIGNALING | 0.12 | 8.44E-22 | 1.41E-20 |
| GLYCOLYSIS | 0.10 | 1.19E-16 | 1.49E-15 |
| APOPTOSIS | 0.09 | 4.40E-12 | 4.40E-11 |
| TNFA_SIGNALING_VIA_NFKB | 0.08 | 6.40E-12 | 5.33E-11 |
| KRAS_SIGNALING_UP | 0.06 | 1.06E-08 | 6.64E-08 |
| MITOTIC_SPINDLE | 0.06 | 1.06E-08 | 6.64E-08 |
| P53_PATHWAY | 0.06 | 1.06E-07 | 5.91E-07 |
| ANGIOGENESIS | 0.17 | 1.26E-07 | 6.32E-07 |
| HALLMARK_UV_RESPONSE_DN | 0.06 | 5.31E-07 | 2.41E-06 |
| CHOLESTEROL_HOMEOSTASIS | 0.09 | 6.10E-07 | 2.54E-06 |
| G2M_CHECKPOINT | 0.05 | 9.69E-07 | 3.46E-06 |
| INFLAMMATORY_RESPONSE | 0.05 | 9.69E-07 | 3.46E-06 |
| SPERMATOGENESIS | 0.06 | 3.41E-06 | 1.14E-05 |
| E2F_TARGETS | 0.05 | 7.98E-06 | 2.35E-05 |
| IL2_STAT5_SIGNALING | 0.05 | 7.98E-06 | 2.35E-05 |
| ADIPOGENESIS | 0.04 | 5.88E-05 | 1.63E-04 |
| PI3K_AKT_MTOR_SIGNALING | 0.06 | 7.15E-05 | 1.88E-04 |
| FATTY_ACID_METABOLISM | 0.04 | 9.03E-05 | 2.15E-04 |
| UV_RESPONSE_UP | 0.04 | 9.03E-05 | 2.15E-04 |
| ESTROGEN_RESPONSE_LATE | 0.04 | 3.83E-04 | 8.33E-04 |
| MYC_TARGETS_V1 | 0.04 | 3.83E-04 | 8.33E-04 |
| TGF_BETA_SIGNALING | 0.07 | 4.46E-04 | 9.30E-04 |
| ANDROGEN_RESPONSE | 0.05 | 5.61E-04 | 1.12E-03 |
| APICAL_JUNCTION | 0.03 | 2.18E-03 | 3.76E-03 |
| COMPLEMENT | 0.03 | 2.18E-03 | 3.76E-03 |
| MYOGENESIS | 0.03 | 2.18E-03 | 3.76E-03 |
| XENOBIOTIC_METABOLISM | 0.03 | 2.18E-03 | 3.76E-03 |
| COAGULATION | 0.04 | 2.26E-03 | 3.76E-03 |
| PROTEIN_SECRETION | 0.04 | 3.78E-03 | 6.10E-03 |
| BILE_ACID_METABOLISM | 0.04 | 6.52E-03 | 1.02E-02 |
| ALLOGRAFT_REJECTION | 0.03 | 1.06E-02 | 1.52E-02 |
| ESTROGEN_RESPONSE_EARLY | 0.03 | 1.06E-02 | 1.52E-02 |
| HEME_METABOLISM | 0.03 | 1.06E-02 | 1.52E-02 |
| PEROXISOME | 0.03 | 3.15E-02 | 4.37E-02 |

**Supplementary Table S4.** Gene set enrichment analysis of differentially expressed genes in MIR211-overexpressing A375 melanoma xenografts

| Genomic Region[a] | Refseq | Gene | Peak ID[b] | length | p-value | Fold Enrichment[c] | q-value |
|---|---|---|---|---|---|---|---|
| chr17:41843518-41843600 | NM_004090 | DUSP3 | S23_peak_6881 | 172 | 9.76E-11 | 4.6 | 6.75E-07 |
| chr17:41843600-41843689 | NM_004090 | DUSP3 | S21_peak_5357 | 90 | 4.53E-06 | 2.8 | 0.009936 |
| chr17:41845784-41845786 | NM_004090 | DUSP3 | S21_peak_5358 | 108 | 4.98E-06 | 2.2 | 0.010662 |
| chr17:41843753-41843833 | NM_004090 | DUSP3 | S23_peak_6882 | 81 | 3.27E-05 | 2.3 | 0.041924 |
| chr17:41845786-41845829 | NM_004090 | DUSP3 | S23_peak_6883 | 106 | 1.78E-13 | 3.8 | 2.55E-09 |
| chr8:29192622-29192688 | NM_001394 | DUSP4 | S23_peak_14673 | 67 | 2.47E-05 | 1.8 | 0.032562 |
| chr8:29193805-29193863 | NM_001394 | DUSP4 | S23_peak_14674 | 59 | 1.08E-05 | 1.9 | 0.015964 |
| chr8:29197663-29197726 | NM_001394 | DUSP4 | S23_peak_14675 | 64 | 1.70E-06 | 2.4 | 0.003438 |
| chr12:89743111-89743112 | NM_001946 | DUSP6 | S21_peak_3124 | 56 | 2.35E-05 | 1.9 | 0.03739 |
| chr12:89743166-89743175 | NM_001946 | DUSP6 | S23_peak_3976 | 64 | 3.70E-05 | 1.8 | 0.046672 |
| chr12:89745439-89745633 | NM_001946 | DUSP6 | S21_peak_3125 | 195 | 5.60E-12 | 4.6 | 6.97E-08 |
| chr12:89745908-89746077 | NM_001946 | DUSP6 | S23_peak_3977 | 170 | 3.45E-07 | 3.4 | 0.000881 |
| chr2:183962890-183962950 | NM_080876 | DUSP19 | S21_peak_7263 | 61 | 2.22E-07 | 5.1 | 0.000726 |
| chr11:102220458-102220491 | NM_001166 | BIRC2 | S21_peak_2502 | 55 | 4.98E-06 | 2.2 | 0.010662 |
| chr11:102220491-102220512 | NM_001166 | BIRC2 | S21_peak_2502 | 55 | 4.98E-06 | 2.2 | 0.010662 |
| chr11:102221400-102221459 | NM_001166 | BIRC2 | S21_peak_2503 | 60 | 3.34E-11 | 6.6 | 3.48E-07 |

**Supplementary Table S5.** The enrichment of *BIRC2* and DUSPs RNAs by Ago2-immunopurification
[a] hg19 genomic coordination
[b] S21 and S23 are two independent RIP-seq replicates.
[C] Fold enrichment compared to input.

| MIR211 low expressors | | | | | MIR211 high expressors | | | | |
|---|---|---|---|---|---|---|---|---|---|
| Cell line | MIR211 Ct | RNU48 Ct | -ΔCt | AUC vemurafenib resistance | Cell line | MIR211 Ct | RNU48 Ct | -ΔCt | AUC vemurafenib resistance |
| UACC-1097 | 38.4 | 22.9 | -15.44 | 1.05 | UACC-2994 | 27.8 | 21.6 | -6.16 | 1.08 |
| UACC-3291 | 38.3 | 22.9 | -15.41 | 0.49 | UACC-0903 | 28.6 | 23.1 | -5.47 | 0.50 |
| UACC-0558 | 38.4 | 23.0 | -15.37 | 0.72 | SK-MEL-119 | 27.9 | 22.5 | -5.41 | 1.14 |
| A375 | 38.1 | 23.1 | -15.08 | 0.50 | SK-MEL-217 | 27.2 | 22.1 | -5.08 | 1.05 |
| UACC-0502 | 38.3 | 23.4 | -14.94 | 0.68 | UACC-3312 | 25.3 | 21.9 | -3.44 | 1.06 |
| UACC-3337 | 37.3 | 22.7 | -14.63 | 1.01 | SK-MEL-113 | 25.4 | 22.3 | -3.07 | 1.15 |
| UACC-2331 | 36.8 | 22.6 | -14.25 | 0.82 | UACC-1649 | 25.9 | 23.0 | -2.88 | 1.12 |
| UACC-1118 | 37.3 | 23.4 | -13.91 | 1.08 | UACC-0257 | 25.3 | 22.6 | -2.67 | 0.67 |
| SK-MEL-2 | 36.6 | 22.9 | -13.71 | 1.04 | UACC-2427 | 27.7 | 25.7 | -1.99 | 1.05 |
| UACC-1237 | 35.9 | 22.2 | -13.64 | 0.81 | UACC-1120 | 24.0 | 22.1 | -1.92 | 0.65 |
| UACC-2641 | 35.3 | 22.3 | -12.99 | 0.95 | MeWo | 24.2 | 22.5 | -1.69 | 1.09 |
| UACC-1729 | 35.9 | 22.9 | -12.97 | 0.96 | UACC-0091 | 23.7 | 22.8 | -0.96 | 0.73 |
| UACC-1113 | 35.3 | 22.4 | -12.86 | 1.03 | UACC-2972 | 22.9 | 22.1 | -0.83 | 1.24 |
| UACC-0612 | 35.4 | 22.6 | -12.83 | 0.93 | UACC-1308 | 23.0 | 22.6 | -0.38 | 0.48 |
| UACC-0647 | 35.6 | 22.9 | -12.71 | 0.99 | UACC-1940 | 25.6 | 25.5 | -0.09 | 1.20 |
| UACC-3074 | 34.6 | 22.1 | -12.52 | 1.05 | UACC-3093 | 24.3 | 24.4 | 0.08 | 1.01 |
| UACC-1093 | 36.2 | 23.9 | -12.28 | 0.93 | SK-MEL-21 | 22.2 | 22.5 | 0.30 | 1.22 |
| UACC-0952 | 34.9 | 23.3 | -11.60 | 1.01 | | | | | |
| UACC-1469 | 34.7 | 23.2 | -11.55 | 1.04 | | | | | |
| UACC-2496 | 34.8 | 23.6 | -11.19 | 0.94 | | | | | |
| UACC-2851 | 33.5 | 22.9 | -10.58 | 1.00 | | | | | |
| UACC-2610 | 33.1 | 22.5 | -10.55 | 1.04 | | | | | |

**Supplementary Tabel S6.** The level of MIR211 expression and vemurafenib resistance of various melanoma cell lines. (-ΔCt = **-**(Ct of MIR211 – Ct of RNU48) from qRT-PCR)

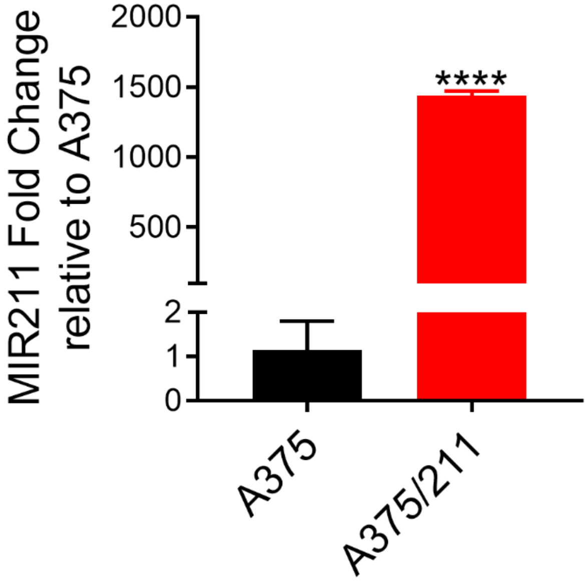


**Supplementary Figure S1.** qRT-PCR-detected changes in MIR211 in A375 and A375/211 cells. (unpaired student t-test, **** $p \leq 0.0001$)

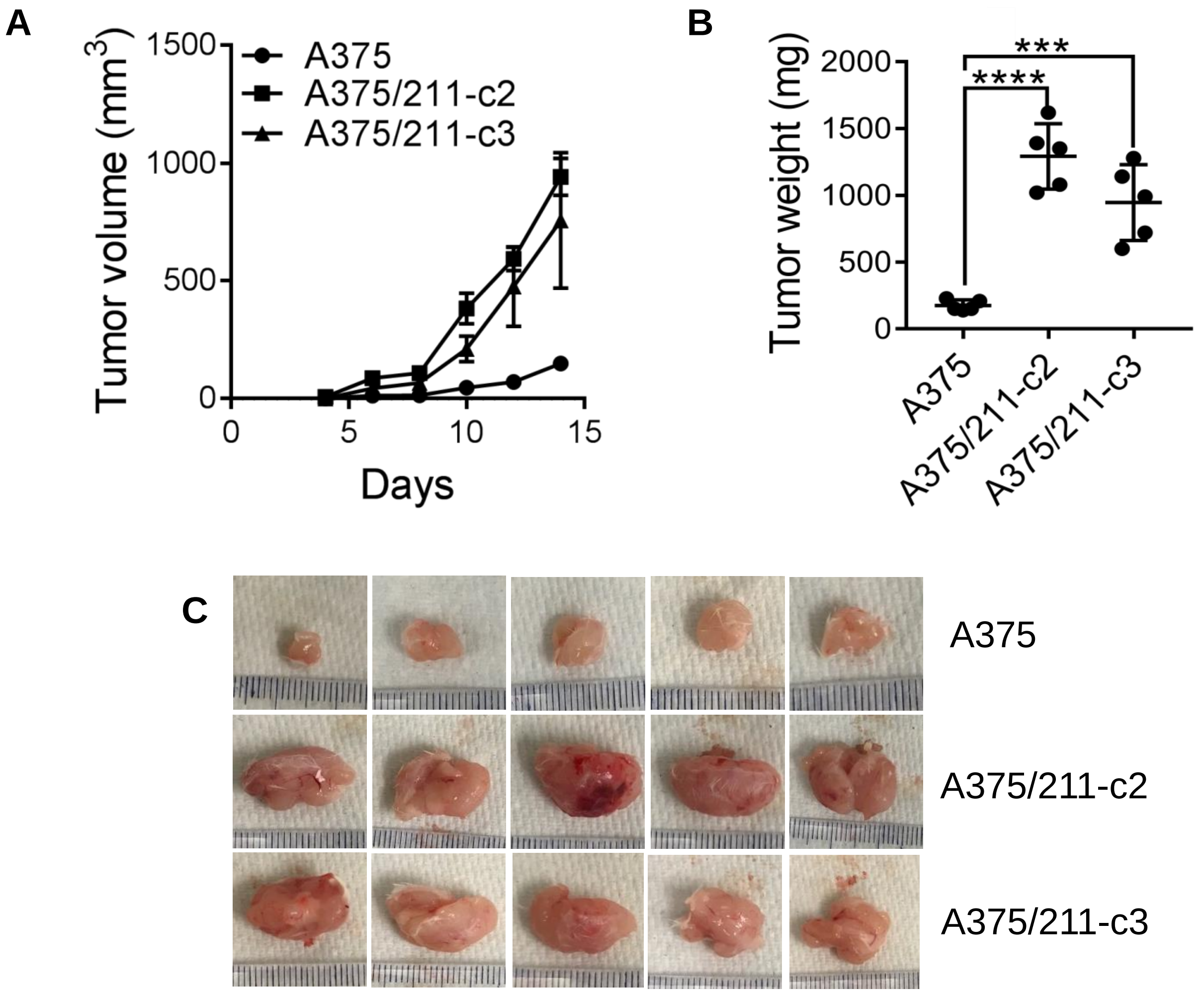


**Supplementary Figure S2.** Different MIR211-overexpressing A375 clones grown as xenografts *in vivo*.

(A) A375 amelanotic melanoma cells and their MIR211 overexpressing counterparts (A375/211-c2 and -c3) were injected subcutaneously into the flanks of SCID mice and tumor volume measured with electronic calipers until termination of the experiment at 12 days. A375/211 tumors were significantly larger and grew more rapidly than the other tumors.

(B) Tumors were excised and weighed at the termination of the experiment at 12 days. MIR211-overexpressing tumors were significantly heavier than either parental A375 xenografts. (unpaired student t-test, *** $p < 0.001$, *** $p < 0.0001$)

(C) Photographs of subcutaneous tumor xenografts at the termination of the experiment at day 12. MIR211-overepxressing xenografts were larger than parental controls.

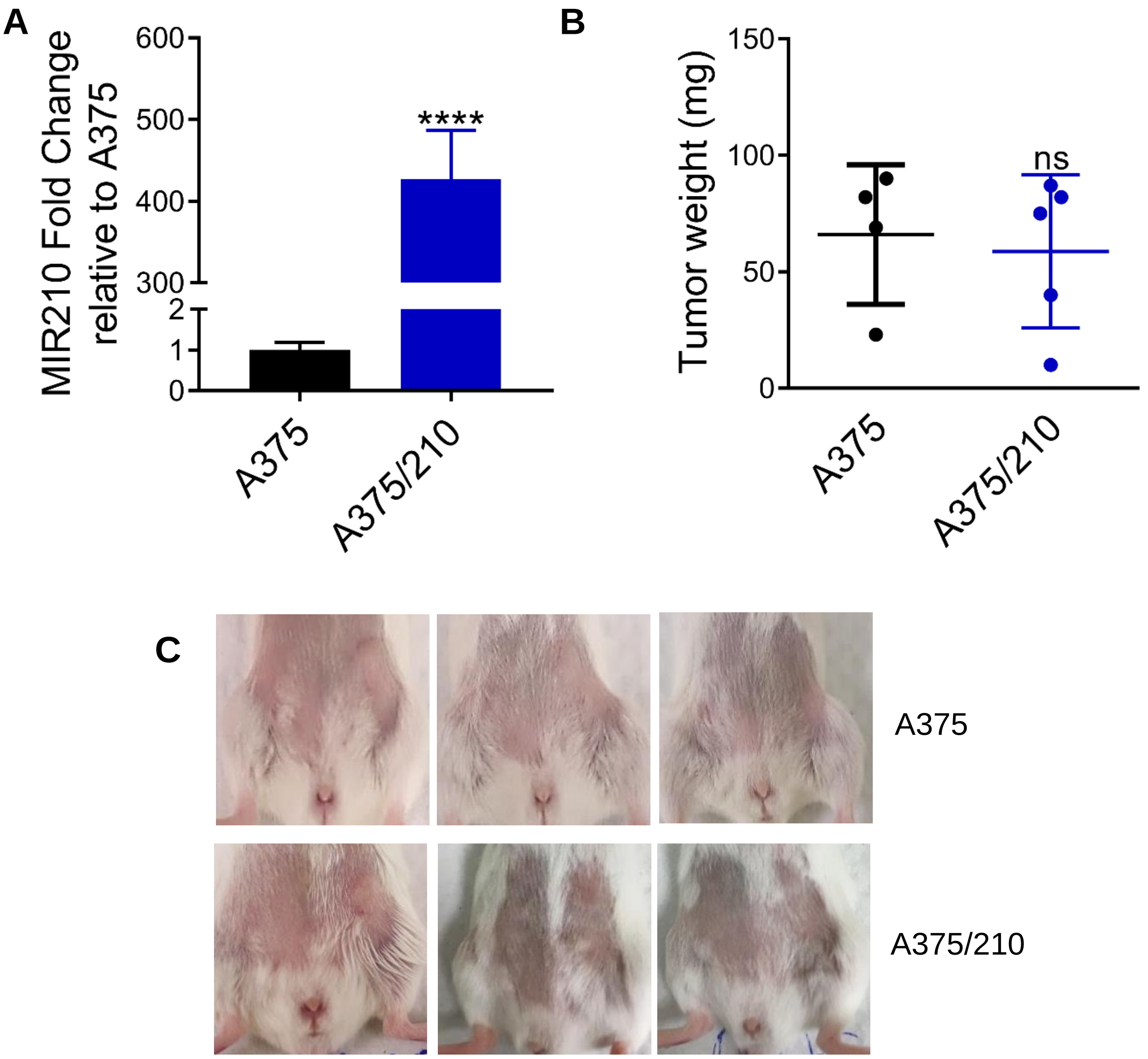


**Supplementary Figure S3.** (A) qRT-PCR-detected changes in MIR210 in A375 and A375/210 cells (B) Tumors were excised and weighed at the termination of the experiment at 12 days. MIR210-overexpressing tumors were not significantly different to parental A375 xenografts. (unpaired student t-test, **** $p \leq 0.0001$, ns: not significant) (C) Photographs of subcutaneous tumor xenografts at the termination of the experiment at day 12.

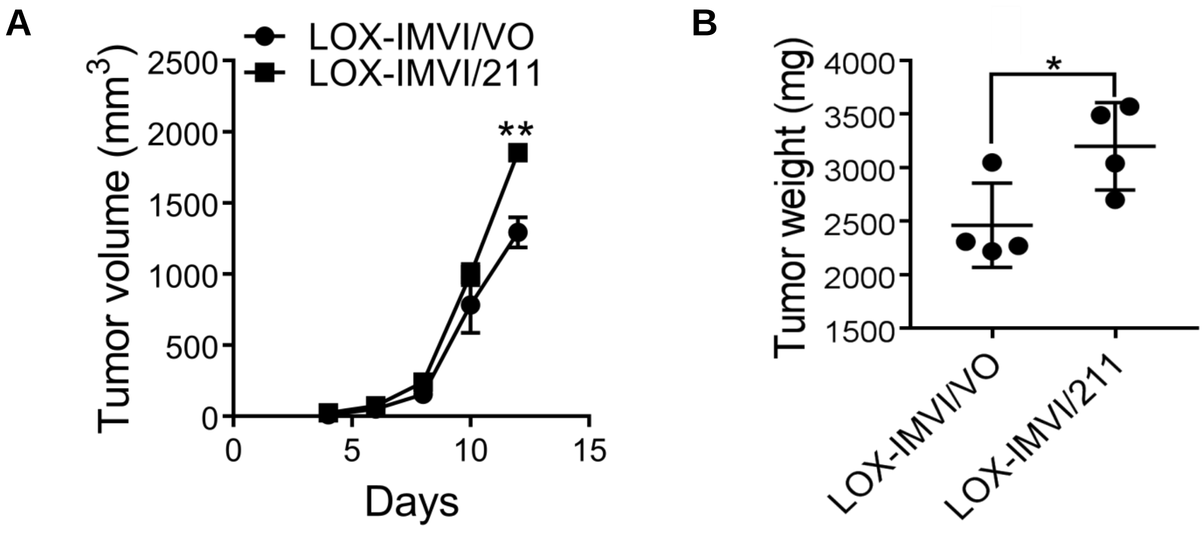


**Supplementary Figure S4.** Ectopic expression of MIR211 in LOX-IMVI cells increased tumor growth. LOX-IMVI stably expressing MIR211 cells were implanted in the flanks of SCID mice. (A) Tumor size was measured and tumor volume was calculated as described in the Methods. (B) Mice were sacrificed at day 12 and tumors were excised and weighed. (LOX-IMVI/VO: LOX-IMVI with vector only; LOX-IMVI/211: cells expressing MIR211, unpaired student's *t*-test, * $p \leq 0.05$, ** $p \leq 0.01$)

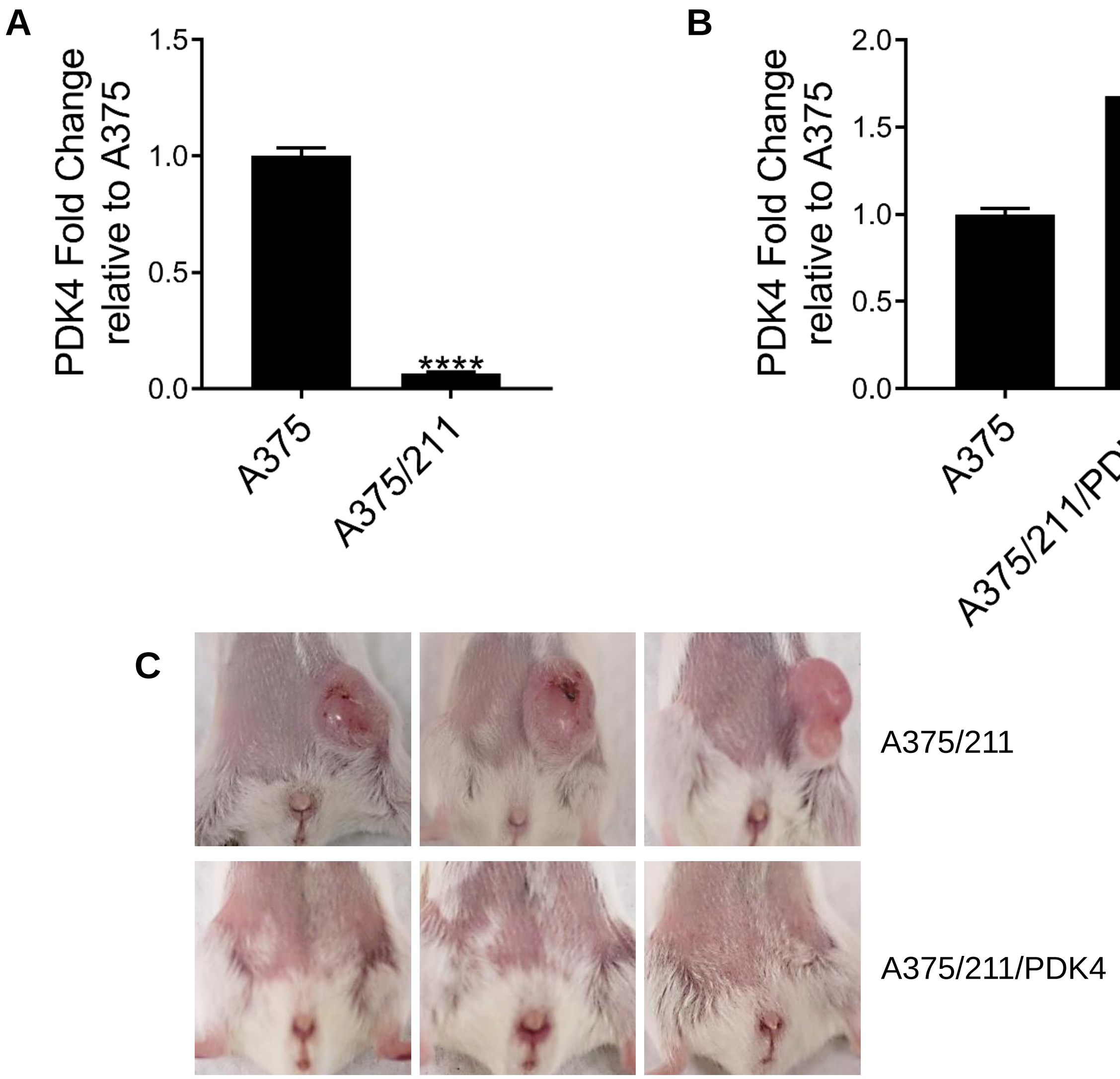


**Supplementary Figure S5.** qRT-PCR-detected changes in PDK4 (A) in A375 and A375/211 cells (B) in A375 and A375/211/PDK4 cells. (unpaired student t-test, ** $p \leq 0.01$ **** $p \leq 0.0001$)
(C) Photographs of subcutaneous tumor xenografts at the termination of the experiment at day 12. The PDK4 expression in A375/211 reduced the tumor growth.

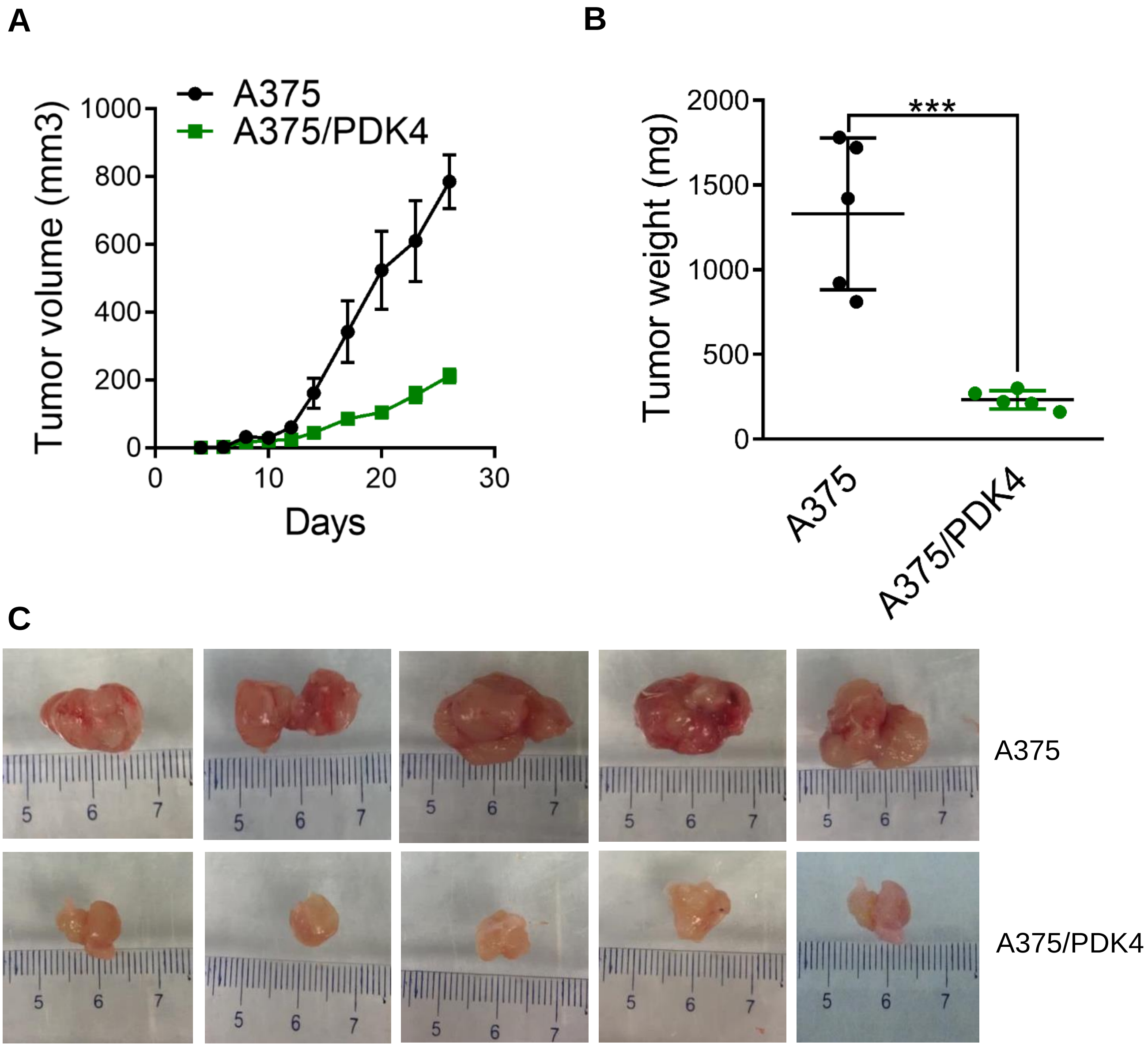


**Supplementary Figure S6.** Ectopic expression of PDK4 reduces tumor growth *in vivo*.

(A) A375 melanoma cells and their PDK4 overexpressing counterparts (A375/PDK4) were injected subcutaneously into the flanks of SCID mice and tumor volume measured with electronic calipers until termination of the experiment at 26 days. A375/PDK4 tumors were significantly smaller and grew more slowly than the other tumors. (student t-test, * $p < 0.05$, ** $p < 0.01$, *** $p < 0.001$)

(B) Tumors were excised and weighed at the termination of the experiment at 26 days. PDK4-overexpressing tumors were significantly lighter than either parental A375 xenografts. (*** $p < 0.001$)

(C) Photographs of subcutaneous tumor xenografts at the termination of the experiment at day 26. PDK4 overepxressing xenografts were smaller than parental controls.

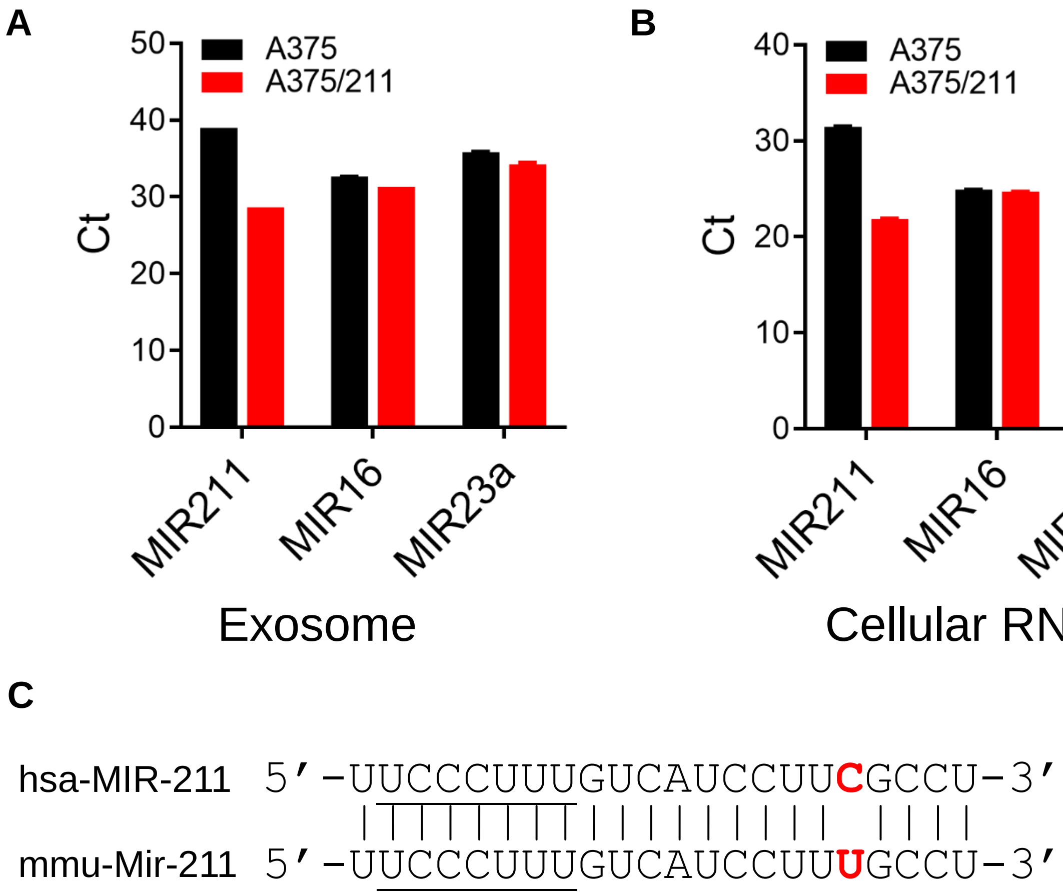


**Supplementary Figure S7.** qRT-PCR showing detection of MIR211, MIR16, and MIR23a (A) in exosomes and (B) cellular RNAs. (C) Differences between the human and mouse MIR211 sequences.

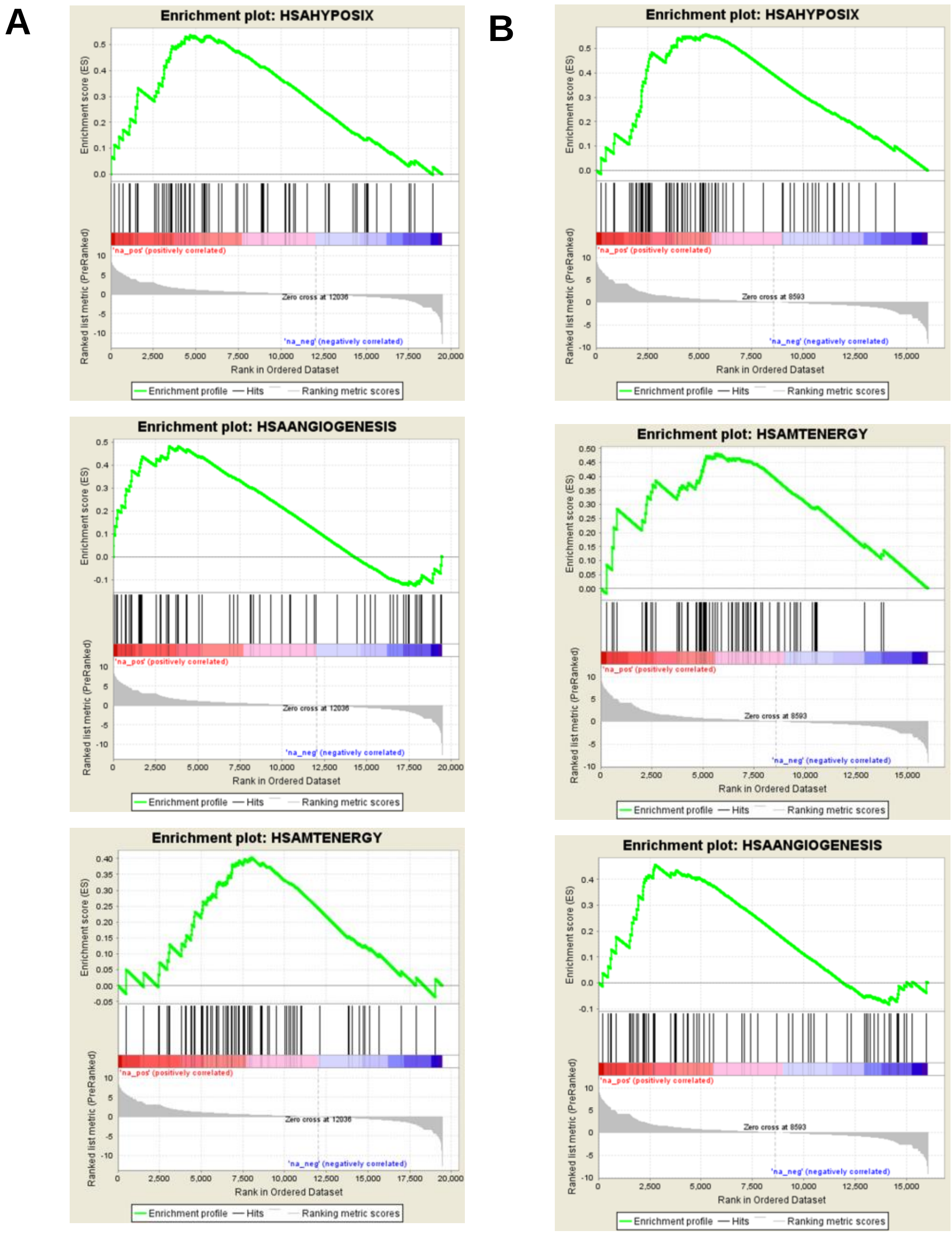


**Supplementary Figure S8.** Gene Set Enrichment Analysis (GSEA) of differentially expressed human (A) and mouse (B) genes in A375/211 vs. A375 xenografts. Hypoxia and angiogenesis pathways were significantly enriched in both human and mouse cells. Mitochondrial Energy metabolism pathways were enriched, but not significantly.

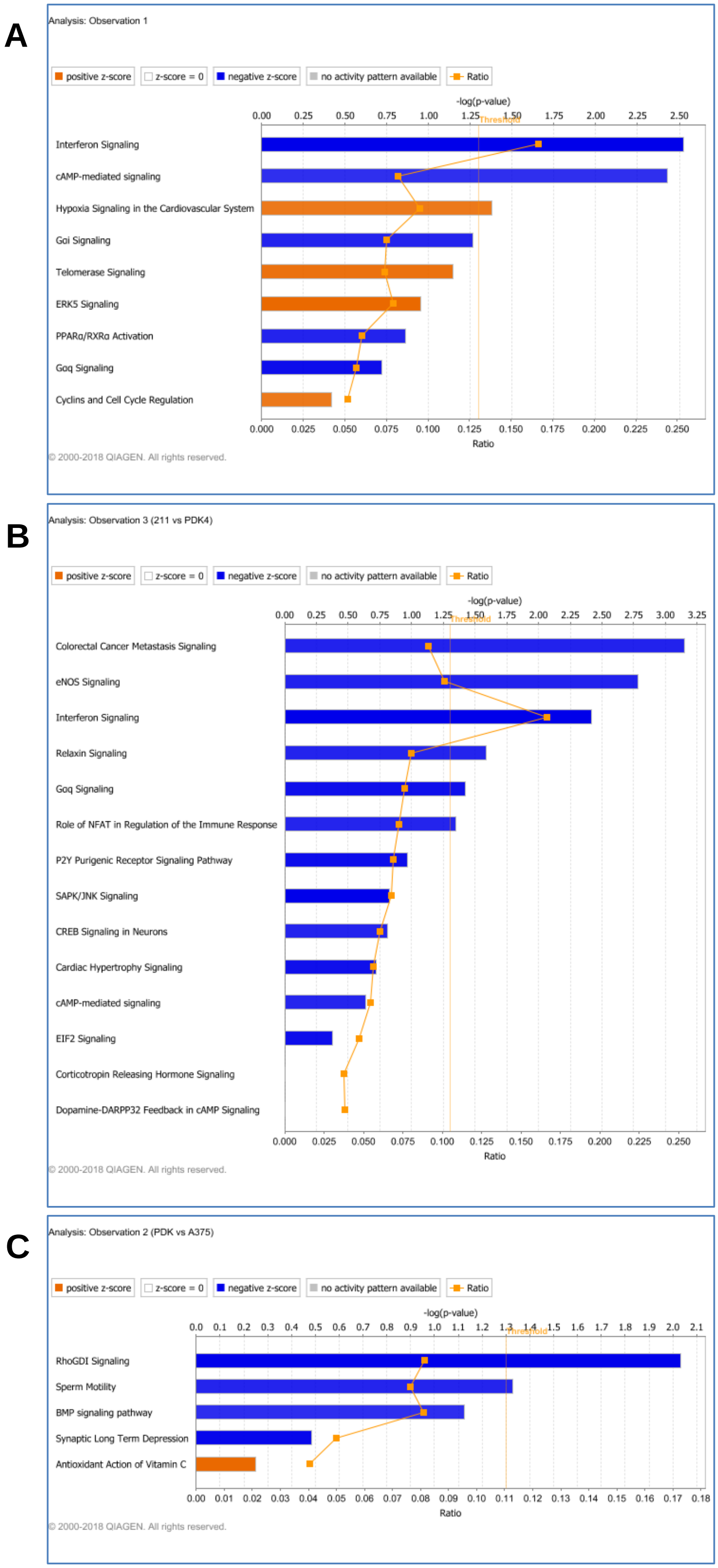


**Supplementary Figure S9**. Canonical Ingenuity Pathway Analysis (IPA) of differential gene expression in (A) A375 vs A375/211 (B) A375/211 vs A375/211/PDK4 and (C) A375/211/PDK vs A375 xenografts.

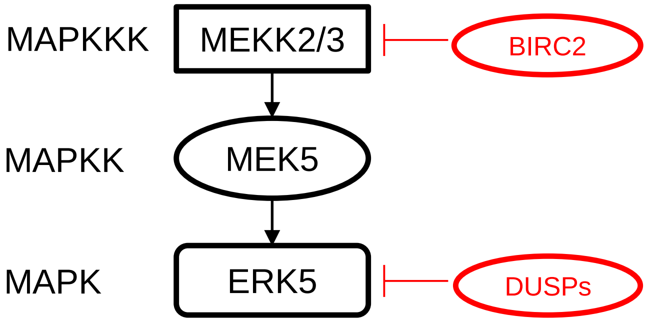


**Supplementary Figure S10**. Schematic diagram of ERK5 pathway. BIRC2 and DUSPs are putative MIR211 targets that negatively regulate ERK5 signaling. BIRC2 inhibits MEKK2/3 by ubiquitination. DUSPs inhibits ERK5 by dephosphorylation.

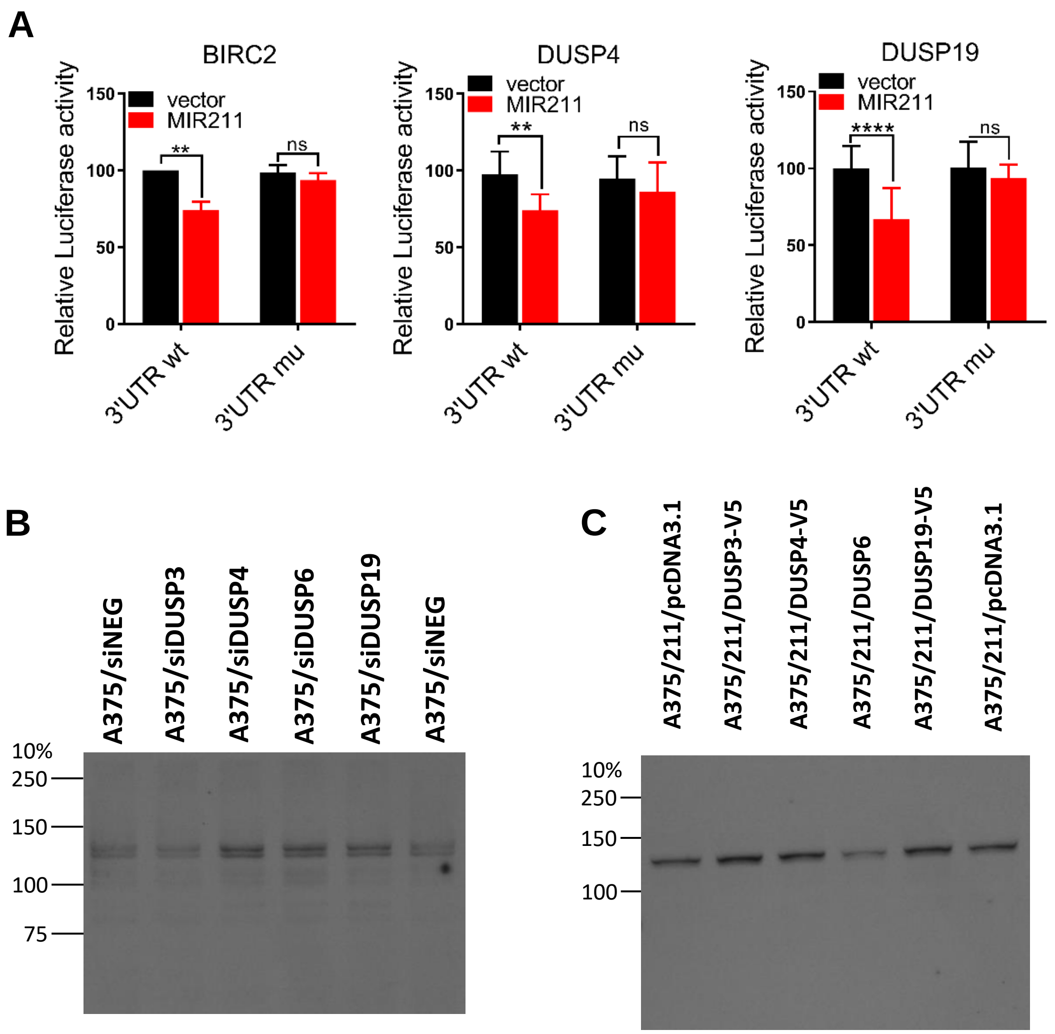


**Supplementary Figure S11**

A. Luciferase reporter assays carrying either wild-type 3'UTR or a mutant with several nucleotides swapped in the region corresponding to the MIR211 seed. When reporter plasmids were co-transfected with plasmids expressing MIR211 into HEK-293T cells, the wild-type BIRC2, DUSP4, and DUSP19 reporters showed reduced luciferase activity, while the mutant one did not. (Student t-test, ns: not significant, ** $p \leq 0.05$, **** $p \leq 0.0001$)

B. phosphor-ERK5 detection by western blot analysis of A375 cells transiently transfected with DUSPs siRNAs.

C. phosphor-ERK5 detection by western blot analysis of A375/211 cells transiently transfected with plasmids harboring each DUSP gene.

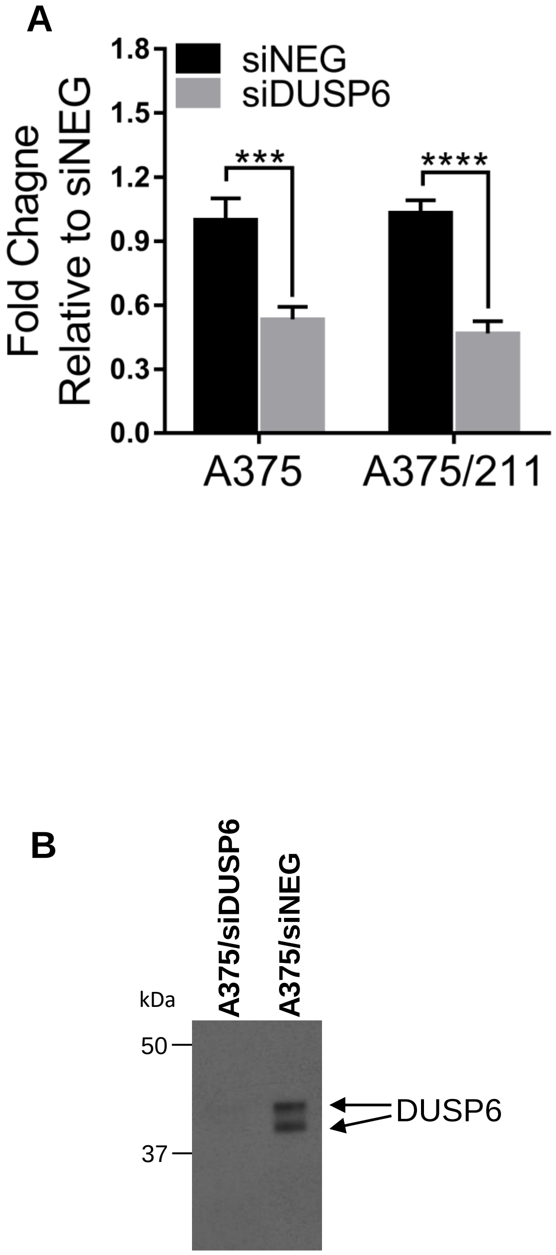


**Supplementary Figure S12.** qRT-PCR (A) and western blot (B) confirmation of DUSP6 knockdown by siRNA.

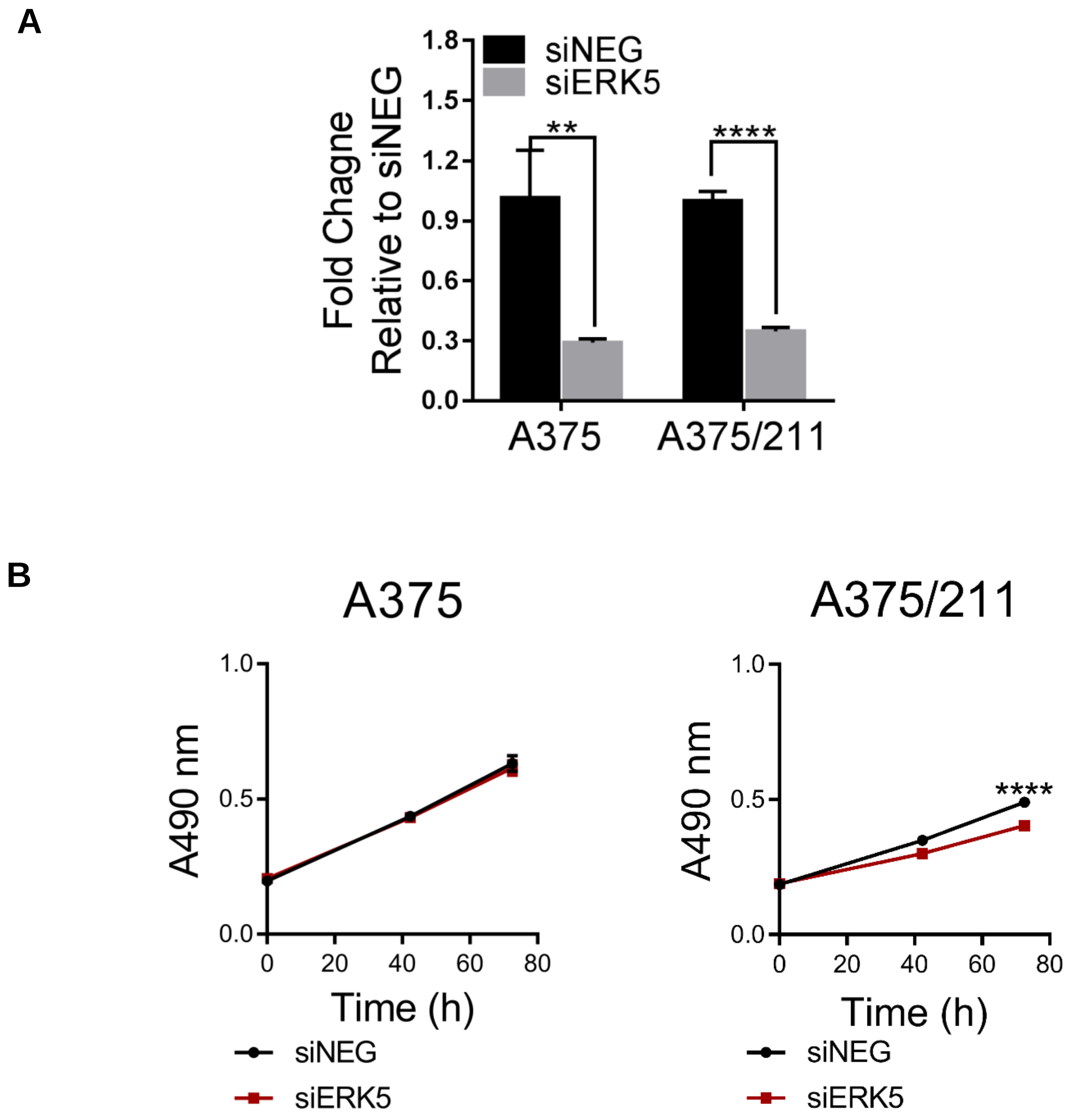


**Supplementary Figure S13.** (A) qRT-PCR confirmation of ERK5 knockdown by siRNA. (B) ERK5 knockdown by siRNA reduced cell viability in A375/211 cells but not A375 cells. (unpaired student t-test, **** $p \leq 0.0001$)

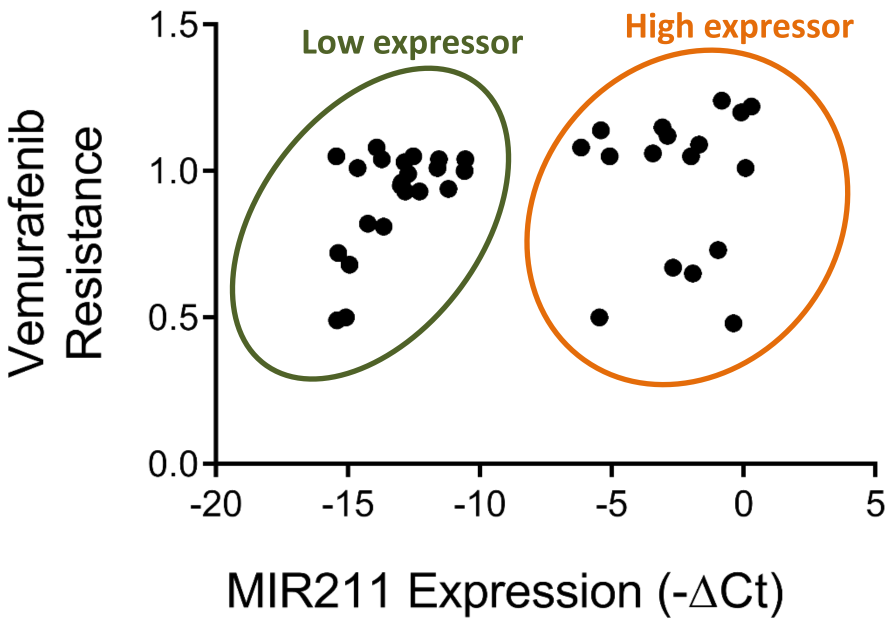


**Supplementary Figure S14.** The level of MIR211 expression and vemurafenib resistance distinguished melanoma cell lines into two groups, MIR211 low expressors and high expressors. Low expressors showed the correlation between MIR211 expression and vemurafenib resistance, while high expressors did not.

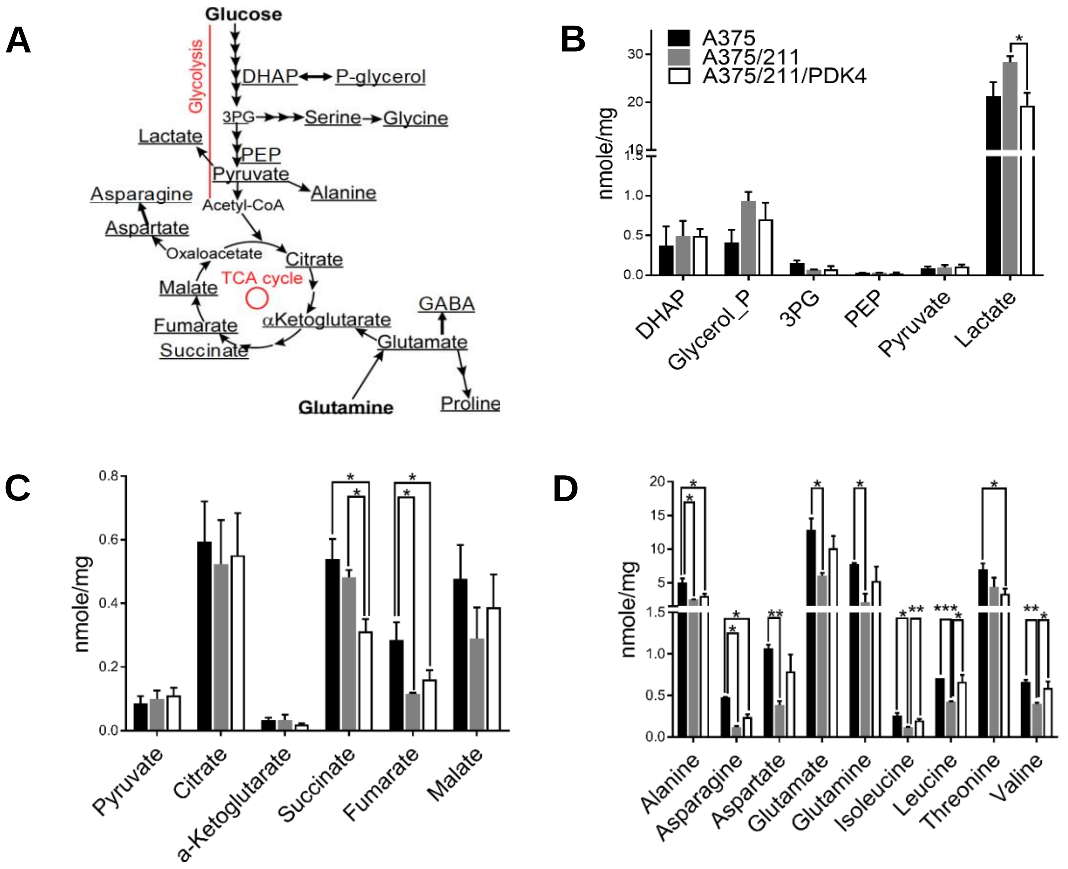


**Supplementary Figure S15.** Targeted metabolomic analysis of A375/211 xenografts.
(A) Schematic of the glycolytic and tricarboxylic cycle (TCA) pathways.
(B-D) *In vivo* targeted metabolomics of (B) glycolysis and glycolytic intermediates, (C) the TCA cycle, and (D) amino acids of A375 (black bars), A375/211 (gray bars), and A375/211/PDK4 (white bars) xenografts. (unpaired student t-test, ** $p \leq 0.05$; ** $p \leq 0.01$, *** $p \leq 0.001$)
